# Affective reactivity during adolescence: Associations with age, puberty and testosterone

**DOI:** 10.1101/524033

**Authors:** Nandita Vijayakumar, Jennifer H. Pfeifer, John C. Flournoy, Leanna M. Hernandez, Mirella Dapretto

**Author notes:** Corresponding Author: Nandita Vijayakumar, School of Psychology, Deakin University, 221 Burwood Hwy, Burwood, VIC, Australia.

## Abstract

Adolescence is a period of heightened social engagement that is accompanied by normative changes in neural reactivity to affective stimuli. It is also a period of concurrent endocrine and physical changes associated with puberty. A growing body of research suggests that hormonal shifts during adolescence impact brain development, but minimal research in humans has examined the relationship between intra-individual changes in puberty and brain function. The current study examines linear and nonlinear changes in affective reactivity in a longitudinal sample of 82 adolescents who underwent three fMRI sessions between the ages of 9 and 18 years. Changes in response to affective facial stimuli were related to age, pubertal stage, and testosterone levels. Using multilevel modelling, we highlight extensive nonlinear development of socio-emotional responsivity across the brain. Results include mid-pubertal peaks in amygdala and hippocampus response to fearful expressions, as well as sex differences in regions subserving social and self-evaluative processes. However, testosterone levels exhibited inverse patterns of association with neural response compared to pubertal stage in females (e.g. U-shaped relationship with the amygdala and hippocampus). Findings highlight potentially unique roles of age, pubertal stage and testosterone on socio-emotional development during adolescence, as well as sex differences in these associations.

---

Adolescence is characterized by shifting social motivations that are accompanied by heightened emotional reactivity and increased sensitivity to social interactions. It is also a period of concurrent biological maturation, including endocrine and physical changes associated with puberty. Animal research indicates that hormonal shifts during this period underlie development of brain regions that subserve social and emotional processes (1, 2). However, longitudinal research with adequate power to detect these developmental processes is limited in humans. Predominant theories of adolescent neurodevelopment historically propose that pubertal maturation drives alterations in subcortical processing of affective stimuli around early adolescence, while age- or experience-related maturation is responsible for more protracted changes in frontal brain regions important for cognitive regulation (3–5). As such, characterizing pubertal associations with brain development also provides an important test of these heuristic “mismatch” models of adolescent neurodevelopment.

## Trajectories of socio-emotional responsivity

One line of investigation on adolescent socio-emotional functioning has focused on the ability to understand the emotional states of others - a crucial skill for successful social interactions. Facial expressions are an important medium for emotional communication, and consequently, research has commonly examined changes in neural responses to affective facial stimuli during adolescence. Early studies revealed greater subcortical activation in the amygdala when passively viewing fearful faces in adolescents compared to adults (6, 7), while others have shown decreased activation with age over this period (8–10). Age-related increases in nucleus accumbens (NAcc) response to happy faces between early and mid-adolescence (11) also suggest there might be differential processing of both negatively- and positively-valenced social stimuli. There is also some evidence supporting adolescent “peaks” in limbic reactivity to affective facial stimuli (primarily using emotional control paradigms) compared to children and adults (12, 13), although studies examining neural responses when passively observing affective facial stimuli have largely failed to identify such non-linear trends (6, 14).

Beyond limbic changes in socio-emotional responsivity, a number of age-related changes have been reported in the prefrontal cortex (PFC), including linear increases in lateral PFC response to fear faces over adolescence (8, 15) as well as a nonlinear mid-adolescent “peak” to affective facial expressions when considering longer age spans (14). Comparatively, converging evidence supports age-related reductions in neural response to affective facial expressions in the medial PFC, including the anterior cingulate cortex [ACC] (7, 16), orbitofrontal cortex [OFC] (7, 14) and ventromedial [vm]PFC (14, 17), although increases may be present when considering early adolescence alone (11). While many of these studies conducted region-of-interest analyses that focused on the PFC, others using a whole brain approach have additionally noted developmental changes in responsivity of the insula, posterior superior temporal sulcus, temporal pole, and fusiform cortices (10, 11, 18–20). In summary, there appears to be widespread changes across limbic, prefrontal, temporal and occipital cortices to affective facial stimuli over adolescence, which leads to the question of what biological processes may be driving these developmental changes.

## Pubertal associations with socio-emotional processes

Following on from research on age-related development, a smaller number of studies have investigated pubertal associations with neural processing of affective facial stimuli. Many of these studies have focused on limbic structures, and in particular the amygdala, given the prevalence of sex hormone receptors in the subcortex (21). Animal research has also shown pubertal changes in neurons and supporting processes, such as pruning of dendrites and synapses, in the amygdala and hippocampus (22–25). Human research using cross-sectional designs have identified reductions in amygdala activation to emotionally neutral facial stimuli with pubertal stage (26) as well as pubertal timing (i.e., pubertal stage relative to same-aged peers, 27). Others have noted reduced striatal activation to fearful and angry faces with earlier pubertal timing (28). However, longitudinal studies have shown increased amygdala and hippocampus response with pubertal stage for both positively- and negatively-valenced emotions (29), as well as increased amygdala and NAcc response for threat-related processes with rising testosterone levels (30). Taken together, there appears to be some inconsistencies in limbic changes identified by cross-sectional and longitudinal studies. Moreover, despite specific hypotheses about puberty-driven peaks in functional reactivity of subcortical structures to affective stimuli (3–5), there have been no longitudinal empirical investigations of nonlinear changes in limbic function elicited by affective faces in relation to pubertal maturation.

Even less is known about puberty-related changes in affective reactivity across the cortex. Cross-sectional findings suggest that adolescents in later pubertal stages (controlling for age) exhibit less ventrolateral (vl)PFC activation to fearful faces, but more activation to angry faces (27). Comparatively, a prior investigation of (a subsample of) the current dataset found increased vlPFC responses to fearful, happy and neutral expressions at later pubertal stages in 13 year olds, as well as increased dorsomedial (dm)PFC response to happy facial expressions, and temporal pole response to happy, sad, fearful and neutral expressions (29). A generalized pattern of increased engagement of occipital cortices to emotional expressions at later pubertal stages has also been reported (29, 31). Based on these limited studies, it appears that pubertal effects may lie within similar regions in the frontal, occipital and temporal cortices that exhibit age-related changes. However, further research is needed to corroborate the pattern of cortical changes identified in this limited literature, as well as better understand the role of pubertal hormones in these changes.

## Current study

The current study aimed to extend the literature on pubertal associations with neural processing of affective facial stimuli, using a single-cohort longitudinal sample that spanned 9 to 18 years of age. While age- and puberty-related trajectories have previously been examined in this typically developing cohort (11, 29), the current investigation incorporated an additional (third) wave of assessments that enabled the modelling of nonlinear developmental trajectories, and also conducted novel analyses involving pubertal hormones. At each assessment, adolescents completed an affective faces fMRI task, where they passively observed exemplars of five different basic emotional expressions in a rapid event-related design. They also completed a questionnaire assessing their pubertal stage and provided a sample of their saliva that was assayed for testosterone levels.

Using multilevel modelling, we investigated linear and quadratic trajectories of subcortical and cortical activation to the affective faces task, using both age and pubertal stage as indices of maturation. We specifically tested for the presence of mid-pubertal “peaks” (i.e., inverted-U shaped trajectories) in neural response to affective facial stimuli. We hypothesized subcortical changes in the amygdala, hippocampus and NAcc, as well as cortical changes in the prefrontal, temporal and occipital regions. However, it was difficult to make hypotheses with further specificity or directionality based on prior research. We also examined the relationship between neural activity and testosterone levels, hypothesizing that hormones may partly underlie puberty-related changes in neural responsivity. Across all analyses, differences in sex and emotional condition were examined. Although these were also largely exploratory in nature, we did speculate that females may exhibit greater adolescent peaks in subcortical and medial PFC activation given they have more intense subjective experiences of, and greater biological reactivity to, affective stimuli compared to their male counterparts during this period (32–36).

## Methodology

### Participants

The sample consisted of 90 adolescents (45 females) aged 9 – 18 years who participated in a three-wave longitudinal study conducted at UCLA. During the first wave (wave 1), participants were aged 9-11 years (M = 10.08, SD = 0.31). Two follow-up assessments were conducted with an approximately 3-year interval between each wave of data collection (see Figure S1). Age ranges and sex distribution at each time point are reported in Table 1. In total, there were 193 (93 male) observations across the three waves, comprised of 33 (18 male) adolescents who participated in one wave of data collection, 11 (6 males) who participated in any (and only) two waves and 46 (21 males) who participated in all three waves. Adolescents and their parents provided written informed assent/consent according to guidelines specified by the Institutional Review Board at UCLA. Participants had no history of diagnosed psychiatric, neurological, or learning disorders (assessed via phone screen), and parent-reported Child Behavior Checklist (37) scores were below clinical range for all syndrome and broadband scales. The final sample size for each measure (PDS, testosterone, fMRI) is summarized in Table 1, with exclusionary criteria discussed in text. Further demographic information on socioeconomic status, race and ethnicity are presented in Table S1. Attrition analyses did not identify any significant differences at wave 1 in demographic or key variables of interest (i.e. socioeconomic status, ethnicity, age, pubertal stage, testosterone levels and gender) in participants who did vs. did not drop out (see Table S2).

**Table 1.** Demographic characteristics and sample size at each time point

|  | Time Point 1 |  | Time Point 2 |  | Time Point 3 |  |
| --- | --- | --- | --- | --- | --- | --- |
|  | M | F | M | F | M | F |
| Age Mean | 10.08 | 10.07 | 13.08 | 13.03 | 16.60 | 16.47 |
| (SD) | (0.34) | (0.30) | (0.35) | (0.28) | (0.59) | (0.64) |
| Overall N | 45 | 45 | 27 | 30 | 21 | 25 |
| PDS N | 44 | 42 | 26 | 27 | 18 | 20 |
| Testosterone N | 45 | 44 | 27 | 27 | 21 | 24 |
| fMRI N | 37 | 39 | 24 | 24 | 16 | 19 |

NB: fMRI sample refers to participants with available PDS and testosterone data as well.

## Materials

### Pubertal Development

At each wave, pubertal stage was measured using the self-report Pubertal Development Scale (PDS; 38). The PDS consists of five questions that assess height growth, body hair and skin changes, as well as breast development and menarche in females, and facial hair and voice changes in males. Items are scored on a 4-point scale ranging from “no physical changes” to “development seems complete”, with the only exception being the yes/no item on menarche. Chronbach’s alpha was 0.89 in females, and 0.85 in males across the three waves. We utilized the Shirtcliff et al. (39) method to score the PDS into a 5-point scale, in order to approximate Tanner staging. In total, 87 participants with 177 measurements were available for analyses (43 females with 89 measurements).

At each wave, participants also provided a saliva sample during the morning of their assessment. Testosterone levels were assayed by Salimetrics. Assays were run in duplicate, and average intra-assay coefficient of variations (CV; based on hormone concentrations) were 4.5% (SD = 4.0%) and 4.3% (SD = 5.4%) for males and females, respectively. Testosterone levels were calculated as the average of the two assays. Measurements were excluded from analyses if, within each wave and each sex, *i)* intra-assay CV was greater than 20% (N = 1 female) or *ii)* mean testosterone levels were greater than 3SD above the mean (N = 2 females). In addition, saliva samples were not provided by two female participants at waves 2 and 3, respectively. This resulted in a sample of 89 participants available for testosterone analyses, with a total of 188 data points over three waves (44 females with 95 measurements). As (mean) testosterone levels were not normally distributed, log-transformed levels were used for analyses (calculated within males and females separately). Mean concentrations and intra-assay CVs at each wave are presented in Table S3. Intra-class correlations for hormones across the three waves are presented in Table S4.

### fMRI paradigm

At each wave, participants undertook a passive fMRI task that involved observing full-color, whole-face basic emotional displays (angry, fearful, happy, sad and neutral) from the NimStim set (40). Events lasted 2s, with an interstimulus interval of variable (jittered) length ranging from 0.5 to 1.5s (M = 1s), and were presented in counterbalanced orders optimized for efficient detection of contrasts between emotions using a genetic algorithm (41). A total of 96 whole-brain volumes were acquired at each time point, comprising 80 events of the stimuli described above and an additional 16 null events (fixation crosses at eye-level). Of the overall sample of 90 adolescents, 86 were available for fMRI analyses. However, a further 3 of these participants were missing either PDS or testosterone data, and one participant’s MRI data was excluded from all waves for failure to meet motion criteria (see below for further detail). As such, the final fMRI sample consisted of 82 participants with a total of 159 observations (41 females with 82 observations).

### fMRI acquisition and analyses

Data were acquired using a Siemens Allegra 3.0 Tesla MRI scanner. A 2D spin-echo scout (TR = 4000 ms, TE = 40 ms, matrix size = 256 × 256, 4-mm thick, 1-mm gap) was acquired in the sagittal plane to allow prescription of the slices to be obtained in the remaining scans. The functional scan lasted 4 min and 54 s (gradient-echo, TR = 3000 ms, TE = 25 ms, flip angle = 90°, matrix size = 64 × 64, FOV = 20 cm, 36 slices, 1.56-mm in-plane resolution, 3-mm thick). For each participant, a high-resolution structural T2-weighted echo-planar imaging volume (spin-echo, TR = 5000 ms, TE = 33 ms, matrix size 128 by 128, FOV = 20 cm, 36 slices, 1.56-mm in-plane resolution, 3-mm thick) was also acquired coplanar with the functional scan. Stimuli were presented to participants through high-resolution magnet-compatible goggles (Resonance Technology, Inc.).

DICOM images were converted to NIFTI format using MRIConvert (http://lcni.uoregon.edu/∼jolinda/MRIConvert/). Prior to preprocessing the brain images, non-brain tissue was removed using robust skull stripping with the Brain Extraction Tool (BET) in FMRIB’s Software Library (FSL; http://www.fmrib.ox.ac.uk/fsl/). Subsequently, images were preprocessed using SPM12 (Welcome Department of Cognitive Neurology, London, UK; http://www.fil.ion.ucl.ac.uk/spm/). This included realignment and co-registration of each subject’s own high-resolution structural image to a mean of the functional images using a six-parameter rigid body transformation model, and reorientation of all (structural and functional) images to the plane containing the anterior and posterior commissures. Images were then spatially normalized into standard stereotaxic space (Montreal Neurological Institute template) using a 12-parameter affine model along with a non-linear transformation involving cosine basic functions. Functional volumes were resampled to 3mm cubic volumes, and then smoothed using a 6mm full-width at half maximum Gaussian filter.

Motion artifacts were identified using an in-house automated script that evaluates changes in image intensity relative to the mean across all subjects, as well as volume-to-volume changes in Euclidean distance (42). This regressor of no interest was constructed by marking volumes of the following types: (a) volumes with greater than 1mm of motion in Euclidian distance relative to the previous volume, (b) volumes for which the *mean intensity across voxels* was extreme (3 SDs above or 1.5 SDs below) relative to the mean intensity across subjects and runs, and (c) volumes for which the *standard deviation across voxels* was extreme (3 SDs above or below) relative to the mean standard deviation across subjects and runs. Volumes immediately preceding and following marked volumes were also flagged (i.e., “sandwich volumes”). The script flagged volumes for excessive head motion in 48 out of 90 participants at time 1 (mean of 11 volumes per run), 32 out of 57 participants at time 2 (mean of 6 volumes per run), and 11 out of 41 participants at time 3 (mean of 3 volumes per run). Significantly more volumes were flagged in males compared to females at time 1 (t_(50)_ = 2.768, p = 0.008), while no differences existed at time 2 or 3 (p > 0.05). Nine participants’ time 1 data, and two participants’ time 2 data, were excluded because > 20% of their volumes were flagged for excessive head motion. In addition, two participants (one from time point 1 and another from time point 2) were excluded as visual inspection of their fixed effects analyses revealed a clear pattern of motion-related striping that indicated contaminated data.

Fixed-effects (i.e., subject-level) statistical analyses were also conducted in SPM12. For each subject, condition effects were estimated according to the general linear model, using a canonical hemodynamic response function, high-pass filtering (128s), AR(1) and no global scaling. A mask of white matter and CSF (https://canlabweb.colorado.edu/wiki/doku.php/help/core/brain_masks) was subtracted from the group average template, and the resultant image was used to restrict the analytic space. Linear contrasts were constructed to assess comparisons of interest within individual participants at each time point; specifically, each emotional expression was contrasted against a resting baseline (as modelled by previous studies using a subset of this dataset, 11, 29). Neutral facial expressions were not employed as a low-level control given they are socially meaningful and often elicit considerable activity in regions associated with processing affective salience and uncertainty (11, 43). Instead, neutral stimuli were modelled as an emotional expression, thus resulting in five contrasts per individual per time point (i.e., activation to happy, angry, sad, fear, and neutral faces relative to rest). These resulting contrasts were used in ROI and whole-brain group-level analyses, conducted within R (v 3.4.0) and AFNI (v17.1.01; https://afni.nimh.nih.gov/), respectively (see next section for further details). ROI analyses were conducted in *a priori* defined clusters in the left and right hippocampus, amygdala and NAcc. Individual anatomical masks for each of these six regions were created using the Harvard-Oxford subcortical atlas, thresholded at 50%. Mean β (parameter) estimates were extracted for these regions from each participant’s fixed effect contrasts using AFNI 3dmaskave. Intra-class correlations across the three waves for each of these ROIs is presented in Table S4.

## Statistical analyses

Linear mixed modeling (LMM) provides a flexible and powerful statistical framework for the analysis of longitudinal data. It is more powerful for detecting developmental trajectories in longitudinal data (comprising three or more waves of repeated measurements) than more commonly used general linear modeling that does not model the covariance structure of serial measurements appropriately and cannot handle imperfect timing and/or subject dropout (i.e., unbalanced data), in particular those cases with only a single time-point (44). We first describe the model-building approach employed for the examination of both age-puberty associations and age-/puberty-related changes in ROI activation (conducted with the lme4 package, 45) in R (v3.4, 46). As we were unable to employ a similar approach for whole brain analyses, next we present the modified null-hypothesis significance testing approach utilized within currently available neuroimaging software.

### Model building approach

First, group-level **developmental** trajectories were investigated in relation to the index of maturation (i.e., age or pubertal stage), by comparing **linear** (Y = Intercept + d_i_ + β1 (maturation) + e_i_) and **quadratic** effects (Y = Intercept + d_i_ + β_1_ (maturation) + β_2_ (maturation^2^) + ei) to the **null** model (Y = Intercept + d_i_ + e_i_). All models were conducted within each i^th^ subject, with a random intercept (d_i_) to account for the repeated observations per subject. The e_i_ represents the normally distributed residual error term. All continuous variables were grand mean-centered. Next, we investigated whether there were significant individual differences in the effect of maturation by adding a **random slope** for maturation to the developmental model (i.e., d(maturation)_i_). Finally, we examined whether the inclusion of **sex effects** improved model fit. We examined whether a main effect of sex (i.e., quadratic model: Y = Intercept + d_1_ + β_1_ (maturation) + β_2_ (sex) + β_3_ (maturation^2^) + e_i_) and an interaction between sex and maturation (i.e., quadratic model: Y = Intercept + di + β1 (maturation) + β_2_ (sex) + β_3_ (maturation*sex) + β_4_ (maturation^2^*sex) + e_i_) improved model fit. At each of these three steps (herein referred to as “developmental”, “random slopes”, and “sex” model comparisons), a more complex model (e.g., linear over null, or sex interaction over main effect) was chosen if likelihood ratio tests indicated the models were significantly different from one another (p < 0.05) and the AIC indicated better model fit (i.e., value smaller by 2 or more). Importantly, this procedure was used to identify a single best-fitting model, and interpretation of coefficients was limited to this final model (i.e., we did not examine the regression coefficients in other models).

### Pubertal analyses

Given prior findings of cubic trajectories of pubertal development (47, 48), we investigated these models in addition to linear and quadratic developmental patterns. The three-step model building approach described above was undertaken to identify the best-fitting model for *i)* PDS development with age (i.e., PDS = Intercept + d_i_ + β_1_ (age) + e_i_), *ii)* testosterone changes in relation to age (i.e., TEST = Intercept + d_i_ + β_1_ (age) + e_i_) and, *iii)* testosterone changes in relation to PDS (i.e., TEST = Intercept + d_i_ + β_1_ (PDS) + e_i_). While sex interactions in these models would indicate significant differences between males and females in a given developmental trajectory, it is possible that this overall group trajectory would not equate to the best-fitting developmental pattern within each sex. Therefore, planned post-hoc analyses were also conducted within each sex separately.

### ROI analyses

Group-level ROI analyses consisted of six model building procedures run in total - one for each of the ROIs. Developmental trajectories of functional activation to *all* emotional faces were investigated in the same model, with *i)* age and *ii)* PDS as the marker of maturation. Given that all five fixed effects contrasts per subject were incorporated into the same model, the step for examining random slopes was modified to investigate if model fit was improved with the addition of a random slope for maturation (i.e., d(maturation)i: 1 + maturation | subject) and a random slope for condition (i.e., d(maturation)i + d(condition)_i_: 1 + maturation + condition | subject). We also conducted an additional (fourth) model comparison step to investigate whether the inclusion of emotional condition effects (i.e., happy, angry, fear, sad, neutral) improved model fit, including its main effect (i.e., quadratic model: Y = Intercept + d_i_ + β_1_ (maturation) + β_2_ (condition) + β_3_ (maturation^2^), its interaction with maturation (i.e., quadratic model: Y = Intercept + d_i_ + β_1_ (maturation) + β_2_ (condition) + β_3_ (maturation*condition) + β_4_ (maturation^2^*condition) + e_i_), and its interaction with maturation and sex (i.e., quadratic model: Y = Intercept + d_i_ + β_1_ (maturation) + β_2_ (condition) + β_3_ (sex) + β_4_ (maturation*condition) + β5 (maturation*sex) + β_6_ (condition*sex) + β_7_ (maturation^2^) + β_8_ (maturation^2^*sex*condition + e_i_). Planned comparisons were conducted within each emotional condition (repeated per ROI). This involved same the model building procedures (without condition effects), and only contained a random intercept as the dataset was not large enough to incorporate a random slope for age (i.e., models failed to converge). Template scripts are available at https://osf.io/4kb6x/.

We also examined the role of testosterone in neural changes. These analyses were conducted in each sex separately, a strategy employed by prior neuroimaging research on puberty (e.g., 49, 50), given the differing role of testosterone in pubertal maturation within males and females (51, 52). That is, any given concentration of testosterone is likely to have different implications for pubertal maturation in males and females, and as such, it does not seem valid to statistically compare the two sexes. Firstly, within each sex, we identified best-fitting PDS models (using the same model building approach described above) and subsequently examined whether testosterone (linear and quadratic terms) improved model fit beyond PDS. Next, we identified best-fitting testosterone models and subsequently examined whether PDS (linear and quadratic terms) improved model fit beyond testosterone.

Finally, we examined the uniqueness of identified trajectories to each variable of interest. To this end, we re-analysed significant findings while controlling for the other maturational/hormonal variables. For example, we re-ran the modelling building procedure for PDS controlling for i) age and ii) testosterone.

### Whole brain analyses

Whole brain analyses were conducted using AFNI 3dLME, which runs voxel-level fMRI group analyses using a LMM approach. We began by investigating developmental trajectories of functional activation to *all* emotional faces with *i)* age and *ii)* PDS as markers of maturation, as well as sex and emotional condition effects. We were unable to employ a similar model-building approach for whole brain analyses given limitations in current neuroimaging software. As such, we employed null-hypothesis significance testing using a full model consisting of linear and quadratic trajectories, along with all sex and emotion main effects and interactions. The first and second-order polynomial terms (i.e., linear and quadratic) were orthogonalized to enable us to interpret significant linear and quadratic terms within the same full model. Given that the combination of a random intercept and slope for maturation provided the best model fit for all ROI analyses (see Results section), whole brain analyses were similarly run with those two terms to account for individual variability. Functional activation was modeled within each subject at each voxel. Reported results exceed the minimum cluster-size threshold needed for a .025 FWE rate (to account for two models) given a voxel-wise threshold of p = .001, with a cluster extent threshold of 38 identified using AFNI 3dClustSim (v17.1.01; 53). Smoothness estimates entered into 3dClustSim were spatial autocorrelation function (acf) parameters averaged from each individual’s group level model residuals, as calculated by 3dFWHMx (age: 0.476 4.72 11.70; PDS: 0.489 4.76 11.81). Un-thresholded group level statistical parameter maps for contrasts of interest can be accessed at https://neurovault.org/collections/3982/.

Given significant sex differences in age- and PDS-related BOLD response in the ACC (ascertained from whole brain analyses), post-hoc analyses were conducted in an independently derived perigenual [p]ACC mask based on a meta-analysis of affective faces neuroimaging research (54), with a 6mm sphere specifically created around the peak of 4 47 7. Within this ROI, we examined linear and quadratic effects of testosterone levels on BOLD response in the pACC ROI in males and females separately, and also the unique effects of age, pubertal stage and testosterone when controlling for one another (as with the subcortical ROIs).

## Results

### Pubertal analyses

Across the entire sample, the relationship between PDS and age was best explained by a group-level (i.e., fixed effect) cubic trajectory that significantly interacted with sex. Accounting for individual differences in the linear effect of age (i.e., random slope) did not significantly improve model fit. As illustrated in Figure 1, the group-level trajectory was characterized by later and steeper development in males compared to females. Post-hoc analyses within each sex identified a similarly significant cubic trajectory in males, and instead a quadratic trajectory in females. Importantly, we do not interpret the cubic trajectory in males to reflect regression in pubertal stage at any point, but is likely an artifact of the model (i.e., imposition of a cubic trend) that is driven by delayed and steeper pubertal maturation relative to females.

**Figure 1.**
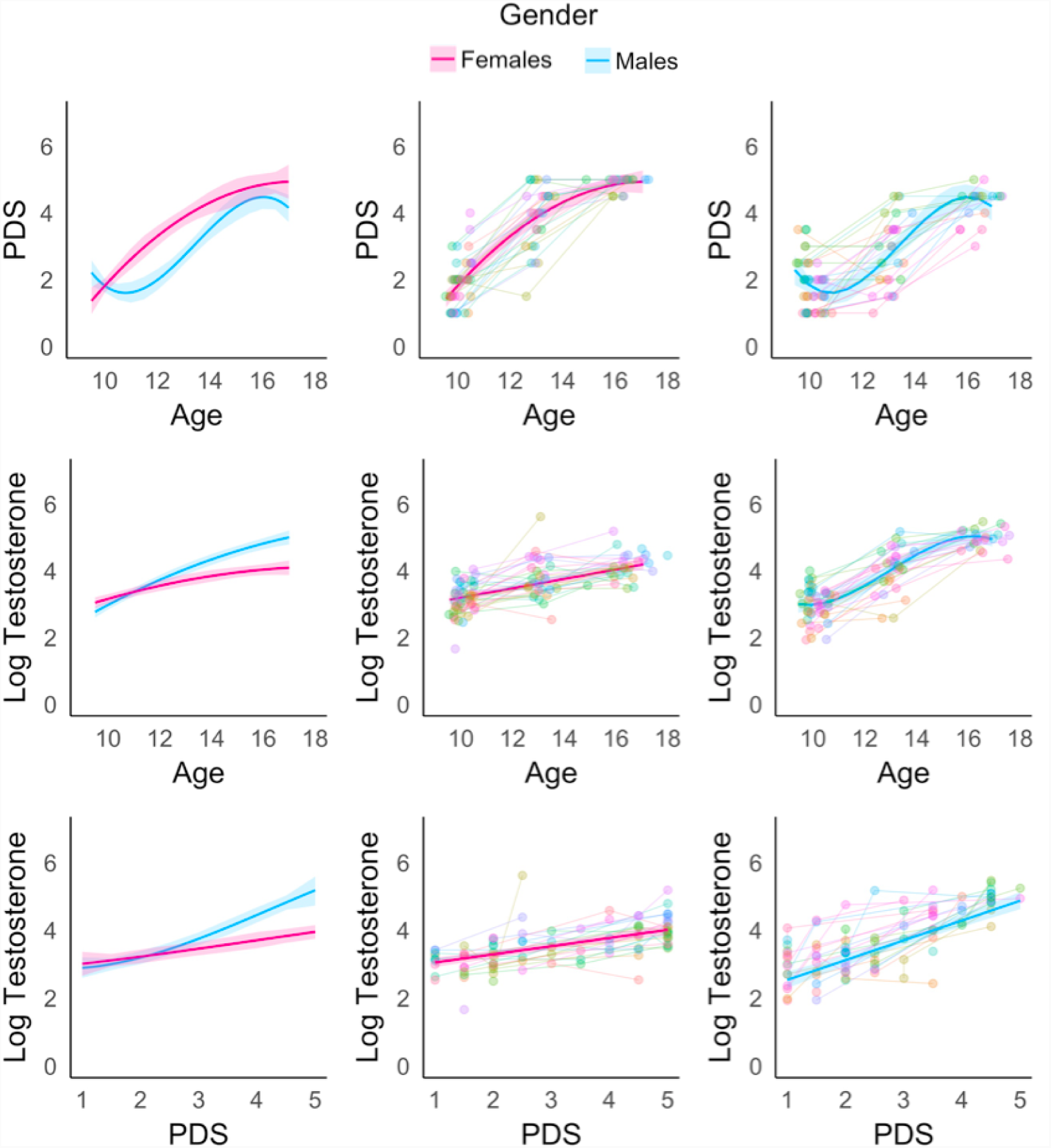
Development of *a)* pubertal stage (PDS) in relation to age, *b)* testosterone (log transformed values) in relation to age, and *c)* testosterone (log transformed values) in relation to PDS. The first column illustrates sex differences in developmental trajectories, while columns two and three illustrate best-fitting developmental models in each sex separately.

The development of testosterone was best explained by a group-level quadratic age-related trajectory that significantly interacted with sex, characterized by greater (nonlinear) increases in testosterone levels within males compared to females (see Figure 1). Again, individual differences in random slope did not improve model fit. Post-hoc analyses within each sex found a significant cubic trajectory in males and linear trajectory in females. In addition, change in testosterone levels was explained by a group-level cubic PDS-related trajectory that significantly interacted with sex, also characterized by greater (nonlinear) increases in testosterone levels with increasing PDS in males compared to females (see Figure 1). Post-hoc analyses within each sex identified significant linear trajectories in both sexes.

## ROI analyses

Across all waves, bilateral amygdala and hippocampus BOLD response significantly increased to affective facial stimuli. Comparatively, the right NAcc exhibited a significant reduction in BOLD response at waves 1 and 3, and the left NAcc exhibited a similar reduction at wave 1 alone (see Table S5).

Results of model fit comparisons are reported in Table S6, and information on random and fixed effects from best-fitting models are reported in Table 2. Corresponding results for models on each emotional condition are reported in Tables S7 and 3.

**Table 2.**
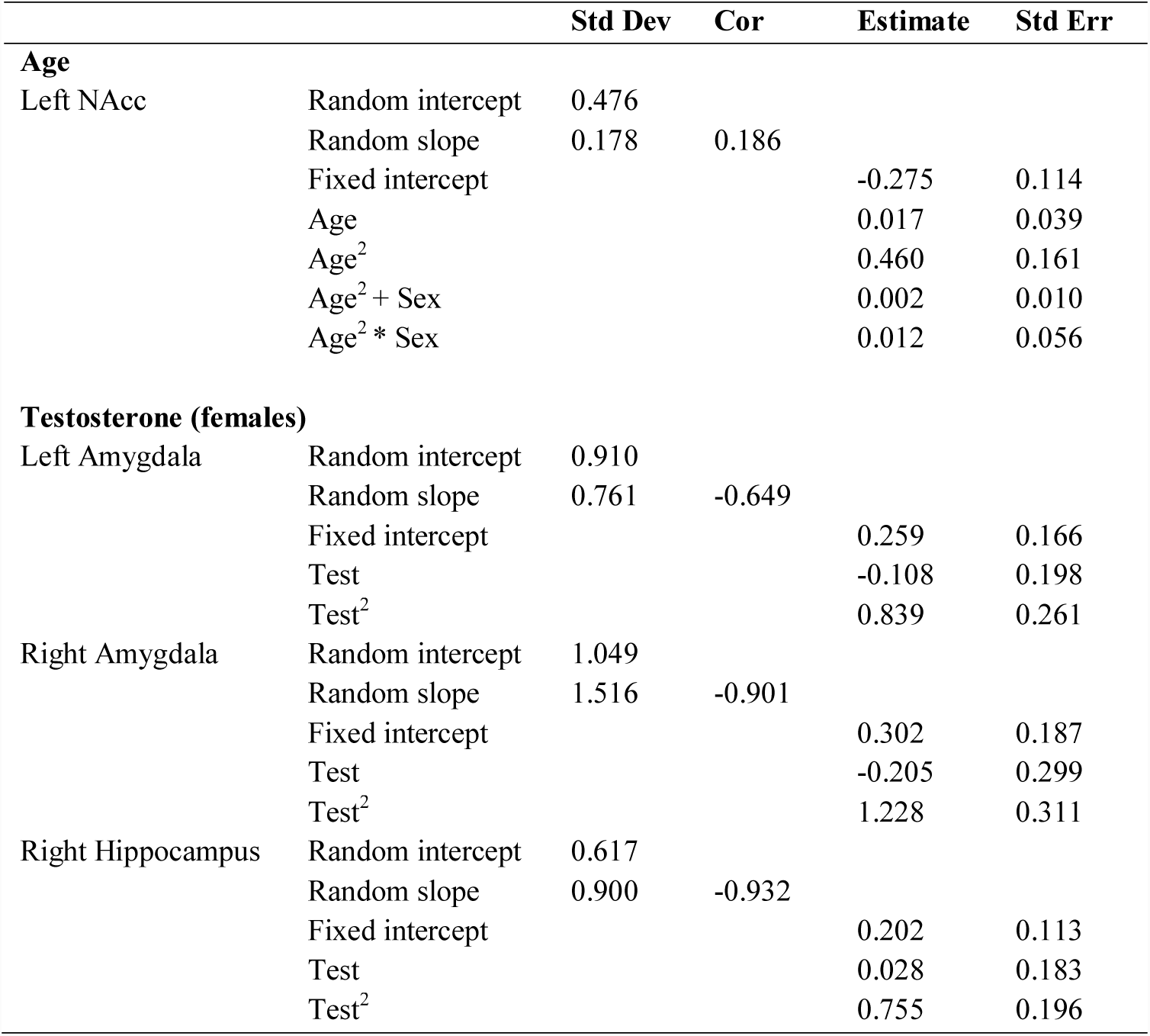
Fixed and random effects of models

|  |  | Std Dev | Cor | Estimate | Std Err |
| --- | --- | --- | --- | --- | --- |
| **Age** |  |  |  |  |  |
| Left NAcc | Random intercept | 0.476 |  |  |  |
|  | Random slope | 0.178 | 0.186 |  |  |
|  | Fixed intercept |  |  | -0.275 | 0.114 |
|  | Age |  |  | 0.017 | 0.039 |
|  | Age<sup>2</sup> |  |  | 0.460 | 0.161 |
|  | Age<sup>2</sup> + Sex |  |  | 0.002 | 0.010 |
|  | Age<sup>2</sup> * Sex |  |  | 0.012 | 0.056 |
| **Testosterone (females)** |  |  |  |  |  |
| Left Amygdala | Random intercept | 0.910 |  |  |  |
|  | Random slope | 0.761 | -0.649 |  |  |
|  | Fixed intercept |  |  | 0.259 | 0.166 |
|  | Test |  |  | -0.108 | 0.198 |
|  | Test<sup>2</sup> |  |  | 0.839 | 0.261 |
| Right Amygdala | Random intercept | 1.049 |  |  |  |
|  | Random slope | 1.516 | -0.901 |  |  |
|  | Fixed intercept |  |  | 0.302 | 0.187 |
|  | Test |  |  | -0.205 | 0.299 |
|  | Test<sup>2</sup> |  |  | 1.228 | 0.311 |
| Right Hippocampus | Random intercept | 0.617 |  |  |  |
|  | Random slope | 0.900 | -0.932 |  |  |
|  | Fixed intercept |  |  | 0.202 | 0.113 |
|  | Test |  |  | 0.028 | 0.183 |
|  | Test<sup>2</sup> |  |  | 0.755 | 0.196 |

**Table 3.** Fixed and random effects of emotion-specific models

|  |  | Std Dev | Estimate | Std Err |
| --- | --- | --- | --- | --- |
| **Age \* Sad** |  |  |  |  |
| Left Hippocampus | Random intercept | 0.182 |  |  |
|  | Fixed intercept |  | 0.632 | 0.124 |
|  | Age |  | 0.083 | 0.034 |
|  | Age<sup>2</sup> |  | -0.034 | 0.015 |
| **Age \* Fear** |  |  |  |  |
| Left NAcc | Random intercept | 0.223 |  |  |
|  | Fixed intercept |  | 0.043 | 0.137 |
|  | Age |  | 0.036 | 0.037 |
|  | Age<sup>2</sup> |  | -0.043 | 0.016 |
| **Fear \* PDS** |  |  |  |  |
| Left Amygdala | Random intercept | 0.528 |  |  |
|  | Fixed intercept |  | 1.006 | 0.184 |
|  | PDS |  | 0.041 | 0.079 |
|  | PDS<sup>2</sup> |  | -0.259 | 0.074 |
| Left Hippocampus | Random intercept | 0.348 |  |  |
|  | Fixed intercept |  | 0.593 | 0.139 |
|  | PDS |  | 0.057 | 0.060 |
|  | PDS<sup>2</sup> |  | -0.169 | 0.057 |
| Right Hippocampus | Random intercept | 0.372 |  |  |
|  | Fixed intercept |  | 0.638 | 0.128 |
|  | PDS |  | 0.046 | 0.055 |
|  | PDS<sup>2</sup> |  | -0.164 | 0.052 |
| **Angry \* PDS** |  |  |  |  |
| Left NAcc | Random intercept | 0.305 |  |  |
|  | Fixed intercept |  | -0.225 | 0.123 |
|  | PDS |  | -0.125 | 0.060 |
|  | Sex |  | 0.453 | 0.178 |
| Left NAcc | Random intercept | 0.414 |  |  |
|  | Fixed intercept |  | -0.122 | 0.089 |
|  | PDS |  | -0.143 | 0.057 |

### Age

Model fit for all six ROIs significantly improved when accounting for individual variability in the random slope of age. BOLD response in the right NAcc was best explained at the group-level by sex-differences in the quadratic trajectory, and post-hoc analyses conducted within each sex revealed that males alone exhibited inverted-U shaped change (p < 0.05). Planned comparisons within each emotional condition found that left hippocampus response during the sad condition and left NAcc response during the fear condition were best explained by group-level quadratic (inverted-U shaped) trajectories. These age-related trajectories are presented in Figure 2. Significant findings remained when controlling for PDS and testosterone (in separate models).

**Figure 2.**
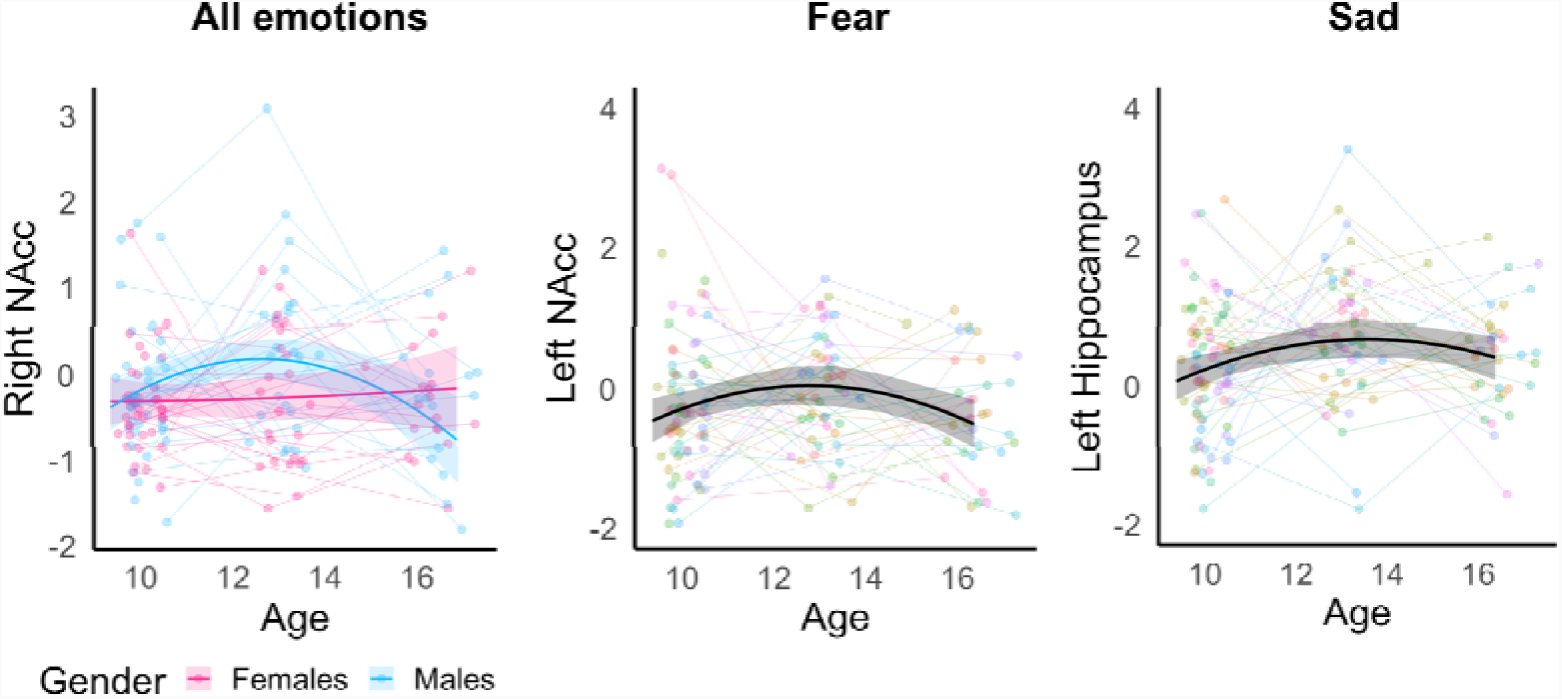
Age-related changes in subcortical response to emotional faces.

### Puberty

Model fit for all six ROIs significantly improved when accounting for individual variability in the random slope of PDS. While none of the subcortical ROIs exhibited significant group-level effects of PDS, planned comparisons within each emotional condition revealed that bilateral hippocampus and left amygdala response during the fear condition were best explained by a quadratic (inverted-U shaped) trajectory (see Figure 3), and bilateral NAcc response during the angry condition was explained by linear reductions. These effects remained when controlling for testosterone and age (in separate models).

**Figure 3.**
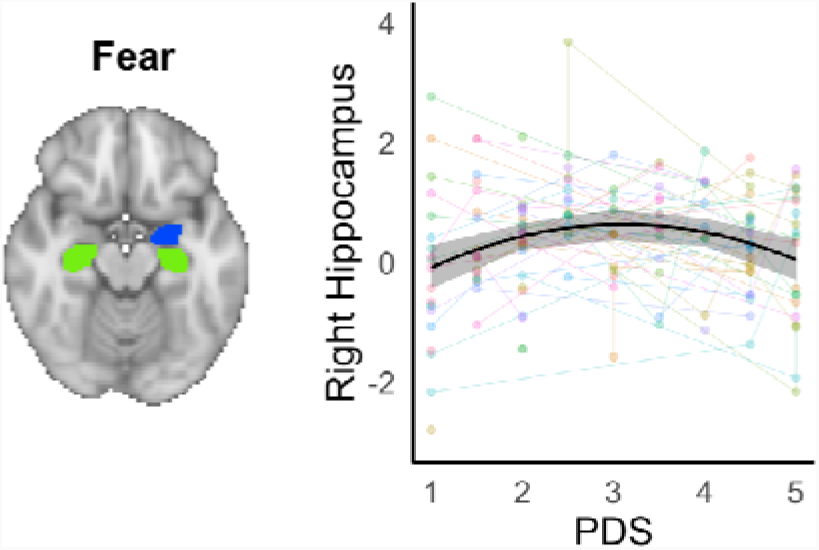
PDS-related quadratic changes in left amygdala and bilateral hippocampus response to fearful expressions. Figure depicts changes in the right hippocampus, but similar patterns were identified in the other two regions (available at https://osf.io/4kb6x/).

### Testosterone

Associations with testosterone levels were examined within each sex separately. In both sexes, across all ROIs, there was significant individual variability in the random (linear) slope of testosterone. In males, none of the ROIs exhibited significant group-level effects. In females, the best-fitting model for the bilateral amygdala and right hippocampus comprised a quadratic (U shaped) group-level association to testosterone levels (as illustrated in Figure 4). These results remained when controlling for age and PDS (in separate models).

**Figure 4.**
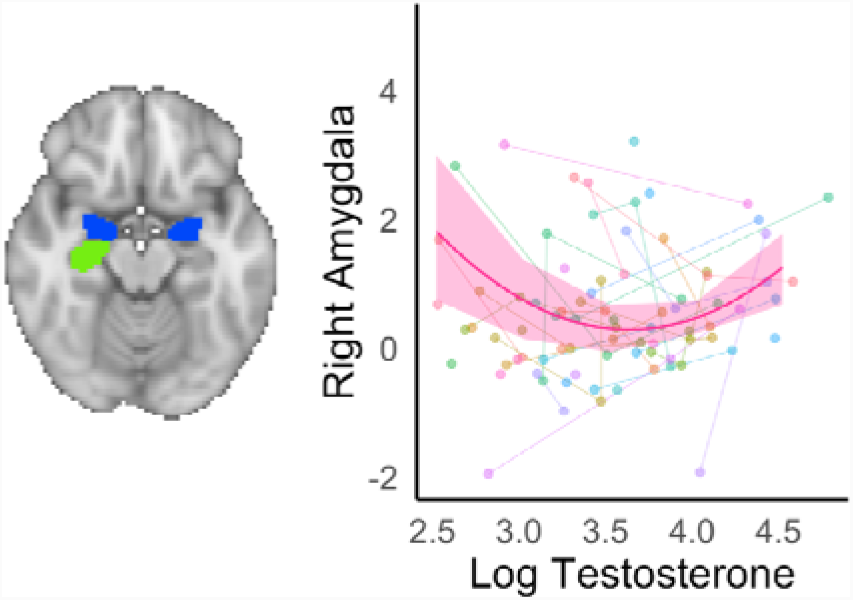
Quadratic changes in bilateral amygdala and right hippocampus response with rising testosterone levels. Figure depicts changes in the right amygdala, but similar patterns were identified in the other two regions (available at https://osf.io/4kb6x/).

### Sensitivity analyses

Employing a stricter motion threshold (maximum of 15% artefact volumes) confirmed all the significant effects for age, pubertal stage and testosterone described above.

## Whole brain analyses

Whole brain analyses of age and PDS did not reveal any significant effects of emotional condition (i.e., main effect or interaction), and the following results represent findings across all five emotional conditions.

### Age

Results of the age model are presented in Table S8. A number of regions exhibited negative quadratic (i.e., inverted U-shaped) trajectories, including multiple clusters of the PFC (i.e., rostral PFC, left ventromedial (vm)PFC and vlPFC), large posterior occipital clusters, left superior parietal and left parahippocampal cortices. Regions exhibiting significant positive quadratic (U-shaped) associations included bilateral precentral and right postcentral cortices, as well as the right lingual gyrus. Sex interactions with quadratic developmental trajectories were also identified within the medial PFC (extending from dmPFC down to ventral ACC and mOFC), right rostrolateral PFC, left mid-cingulate, right parahippocampal, and right anterior temporal cortices (see Figure 5). In these regions, males had inverted U-shaped trajectories, while females had either U-shaped trajectories or minimal change. Opposing patterns of sex interactions with quadratic development (i.e., U-shaped trajectories in males and inverted-U shaped trajectories in females) were identified in multiple occipital and motor cortices, as well as the right OFC.

**Figure 5.**
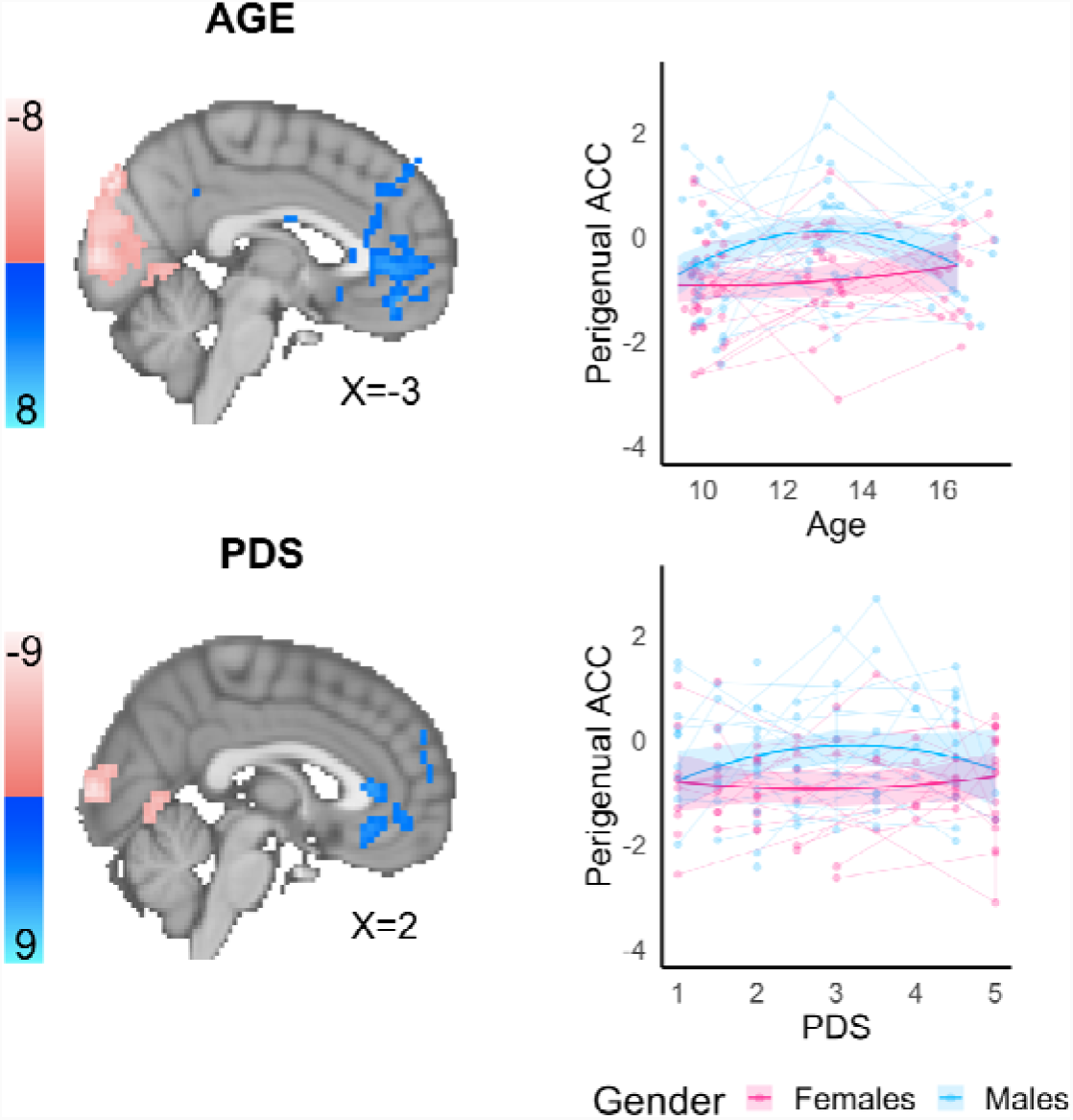
Sex differences in age- and PDS-related quadratic trajectories (Z-scores displayed). Changes in an independently defined perigenual ACC ROI (4 47 7) are plotted.

### Puberty

The PDS model revealed significant linear increases in activation of the lateral occipital regions and the right medial OFC (refer to Table S9). The left temporoparietal junction exhibited significant negative quadratic (inverted U-shaped) associations with pubertal development. In comparison, multiple motor (i.e. supplementary motor, bilateral somatosensory and right premotor) and occipital clusters exhibited positive quadratic (U-shaped) associations. Significant sex interactions with quadratic developmental trajectories were identified within the ventral ACC and vmPFC, right striatum, right dmPFC, and right TPJ (see Figure 5). In these regions, males had inverted U-shaped trajectories, while females had either minimal or U-shaped change. The opposing pattern of sex interactions with quadratic development was identified in the posterior occipital cortices, as well as right dorso/ventrolateral PFC and inferior parietal cortex.

Post-hoc analyses were conducted on an independently defined pACC ROI. Firstly, controlling for testosterone did not change the significance of sex differences in PDS- or age-related trajectories. In separate models, we found that testosterone was not associated with BOLD signal changes in the pACC in males, but rising testosterone levels were associated with significant inverted-U shaped changes in females.

## Discussion

The current study investigated longitudinal changes in functional brain activation to affective facial expressions across early to late adolescence. We identified both linear and nonlinear changes in subcortical response to affective faces, with unique effects of age and self-reported pubertal development. Cortical changes were characterized by sex differences in regions that subserve social cognitive and emotion regulatory processes. Rising testosterone levels, however, did not underlie any of the puberty- or age-related changes.

### Subcortical development

Age-related trajectories in subcortical ROIs included inverted U-shaped changes, or mid-adolescent “peaks” in the left NAcc for fearful expressions, as well as the right NAcc in males for all emotional expressions. A similar pattern was also found in the left hippocampus response to sad faces. While NAcc activation is typically associated with reward sensitivity (55–57), it also responds to salient and/or aversive stimuli and is implicated in successful emotion regulation (58–61). The hippocampal memory system also plays an important role in the regulation of emotional response to negative stimuli, along with its extensive connections to the amygdala and extrastriate visual cortices (62). Thus identified changes in neural sensitivity may reflect age-related changes in engagement of regulatory processes in response to affective stimuli. In particular, the hippocampal findings may reflect social processes that support the recognition of sadness during this period, given the sharp rise in negative affectivity and associated psychopathology between early and mid-adolescence (63). These results also remained significant when controlling for biological changes (i.e. pubertal stage and testosterone levels), which provides some, although not exhaustive, support for the potential role of environmental influences. These findings also highlight the value of examining multiple developmentally appropriate emotional expressions, beyond threat-related processes when considering a chronologically (i.e. socially) defined stage of development.

There has been significant interest in the effect of puberty on limbic reactivity based on animal research showing changes in neurons and supporting processes (i.e. pruning of dendrites and synapses) in these regions during puberty (22–25). Subcortical analyses identified changes in threat-related processes with pubertal maturation, including mid-pubertal “peaks” in the bilateral hippocampus and left amygdala when processing fear expressions and linear reductions in NAcc response when processing angry expressions. Despite hypotheses of pubertal changes in threat-related processes, prior cross-sectional studies have failed to identify changes in limbic response to anger or fear expressions with pubertal maturation (26, 27). While a prior longitudinal analysis of (a subsample of) this dataset revealed puberty-related increases in amygdala and hippocampal response between early and mid-adolescence (29), and the current study extends these findings by highlighting nonlinear change when considering development into late adolescence. Findings remained when controlling for age, suggesting that both progression through the pubertal stages and speed of progression through stages relative to peers (often referred to as pubertal tempo, 64) relate to changes in affective reactivity. Moreover, it appears that puberty and age each have unique effects on fear-related processing, which may suggest a role for both environmental and biological influences.

### Cortical development

Investigation of whole-brain voxel-level data revealed numerous clusters exhibiting nonlinear inverted-U shaped associations with age between early and late adolescence. This effect was identified in the anterior PFC extending to medial OFC, dlPFC, vlPFC, parahippocampus, mid-cingulate and occipital cortices. Overall, this nonlinear pattern may explain some conflicting findings in prior research. For example, a longitudinal analysis of this sample found increased activation of the medial OFC between 10 and 13 years of age (11), but other studies with larger age ranges have found reduced activation into the mid 20s (14, 65). Similarly, studies focusing on late childhood to mid adolescence have identified increased dlPFC and vlPFC responses with age (8, 15), while others noted decreased response when considering the extended span of late childhood to adulthood (18, 65). While others have previously failed to identify nonlinear changes when observing affective faces (e.g. 14) this may reflect the increased power of longitudinal studies to identify developmental trajectories in comparison to cross-sectional studies.

Among the most prominent cortical findings were extensive sex differences in regions subserving social and self-evaluative processes. This included overlapping age- and puberty-related changes across the medial PFC, in addition to puberty-related changes in the TPJ and age-related changes in the left dACC and right anterior temporal cortex. Somewhat similar to the limbic effects in the right NAcc, these regions were all characterized by mid-adolescent peaks in males, versus either mid-adolescent troughs or minimal change in females. While we hypothesized that females would exhibit greater neural reactivity (particularly within regions implicated in socio-emotional processes) given rises in affective (behavioral and physiological) sensitivity relative to male counterparts during early adolescence (34), a prior meta-analysis of adolescent and adult studies found that males exhibited greater subcortical and medial prefrontal reactivity to emotional face stimuli than females (66). Males also show less discrimination in medial PFC activation to positively- and negatively-valenced stimuli, while females display selective activation to unambiguous threatening cues (67). Conversely, sex differences may reflect a potential decoupling of mPFC and limbic response in females, which has been related to increases in withdrawn behaviors during adolescence (68). Further research on coordinated activity of limbic and prefrontal regions, and associations with emotional control, may thus improve our understanding of how sex differences in neurodevelopmental trajectories relate affective behaviors during adolescence.

### Hormonal processes

The prevalence of sex steroid hormone receptors in the brain (21) has led to hypotheses that hormonal shifts during puberty are partly responsible for neurobiological development, including “organizational” and “activational” effects that refer to permanent changes in brain structures and temporary changes in activation within existing neural systems, respectively. However, we found that controlling for testosterone did not alter PDS- (or age-) related trajectories in either the subcortex or cortex. Moreover, in females, rising testosterone levels were associated with opposing patterns of neural changes to PDS, characterized by U-shaped changes in the amygdala and hippocampus, and inverted U-shaped changes in the pACC. These results are somewhat consistent with prior findings of decreased amygdala and increased PFC response with higher testosterone levels during explicit emotion control in mid-adolescents (14 year olds, 69). It has also long been hypothesised that testosterone has fear-reducing properties by decreasing amygdala reactivity to affective stimuli, although empirical support has been mixed (70). However, others have noted increased amygdala response to threatening facial stimuli with rising testosterone levels between early to mid-adolescence (68). As such, it is difficult to interpret our findings in the context of the limited prior literature, but it does appear that testosterone may have unique effects on neural function relative to pubertal stage, consistent with research on structural maturation of these regions (48). It is important that future studies also incorporate assessments of estradiol and DHEA. While these hormones have anxiogenic and depressogenic properties in adults, rodent studies suggest that hormones have differential effects across development (71). As such, a more comprehensive characterization of hormonal changes is needed to better understand potential mechanistic processes during this period.

### Limitations

The current findings should be considered in light of certain limitations. Firstly, testosterone concentrations in saliva samples can be influenced by a number of extraneous factors (e.g., diet, exercise, stress, 72, 73), and future studies should consider collecting multiple saliva samples to calculate basal hormone concentrations. The stimuli in our functional paradigm were comprised of young adult faces given the focus on investigating emotional processing more broadly and limitations in databases available at the time of study conception. However, developmental databases with adolescent stimuli are now freely available, and future investigations may benefit from using age-matched stimuli to investigate peer-related socio-emotional processing given the increased saliva of peer interactions (e.g., 14). Future studies would also benefit from the inclusion of other basic emotions (i.e., surprise and disgust) to improve generalizability of findings that collapse across the difference emotions. Our interpretation of findings as potentially reflecting environmental and/or biological influences are preliminary, given the limited breadth of variables that were examined. A number of factors beyond of scope of this study are known to impact affective reactivity, such as the experience of early life stress (74), and future research that builds on our findings with the incorporation of these variables is needed.

While it is undoubtedly a strength that up to three waves of fMRI data were collected in this study, we can only account for within-person measurement error in longitudinal estimates of linear slopes, and estimates of quadratic effects only account for uncertainty due to variation between participants. We also conducted testosterone analyses in males and females separately, given the differential role of testosterone in each sex (51, 52). Sex differences in brain-hormone associations are thus qualitative and should be interpreted with caution, as significant effects that were identified in one sex alone may not be statistically different from the other sex if directly compared. We did not correct for multiple comparisons for subcortical analyses, as we use a likelihood-based approach that provides information about the relative usefulness of a model to describe data in comparison to another model. It does not, however, provide information about the absolute worth of any given model. Thus the results should be interpreted in terms of relative evidence as opposed to absolute conclusions (i.e., without reference to the comparison models, 75). Finally, examination of individual PDS trajectories in Figure 1 reveals that females were, overall, more pubertally advanced than males at wave 1, and while most females had completed maturation by wave 3, significant variability remained in males. As such, we may have failed to capture pubertal influences on brain function occurring during earlier phases in females, and future studies examining sex differences would benefit from recruiting females at earlier ages relative to their male counterparts (e.g., 30, 68, 76).

### Conclusions

Despite these limitations, this investigation is the first longitudinal functional neuroimaging study of puberty-related socio-emotional development with three waves of data collection. Using multilevel modelling, we identified extensive nonlinear development of socio-emotional functioning across the brain. Amygdala and hippocampus reactivity to fear expressions were characterized by inverted-U shaped changes with pubertal maturation. Sex differences in development were prevalent in regions subserving social and self-evaluative processes. Unexpectedly, testosterone did not appear to underlie puberty-related changes in limbic or prefrontal cortices, with findings overall highlighting unique roles of age, pubertal stage and testosterone on socio-emotional development during adolescence.

## Funding & acknowledgements

NV and JHP were supported by MH107418 (PI: Pfeifer). The project described was supported by Grant Numbers RR12169, RR13642 and RR00865 from the National Center for Research Resources (NCRR), a component of the National Institutes of Health (NIH). For generous support the authors also wish to thank the Santa Fe Institute Consortium, Brain Mapping Medical Research Organization, Brain Mapping Support Foundation, Pierson-Lovelace Foundation, The Ahmanson Foundation, William M. and Linda R. Dietel Philanthropic Fund at the Northern Piedmont Community Foundation, Tamkin Foundation, Jennifer Jones-Simon Foundation, Capital Group Companies Charitable Foundation, Robson Family and Northstar Fund.

