## Supplementary material for "Affective reactivity during adolescence: Associations with age, puberty and testosterone"

**Supplementary Information**

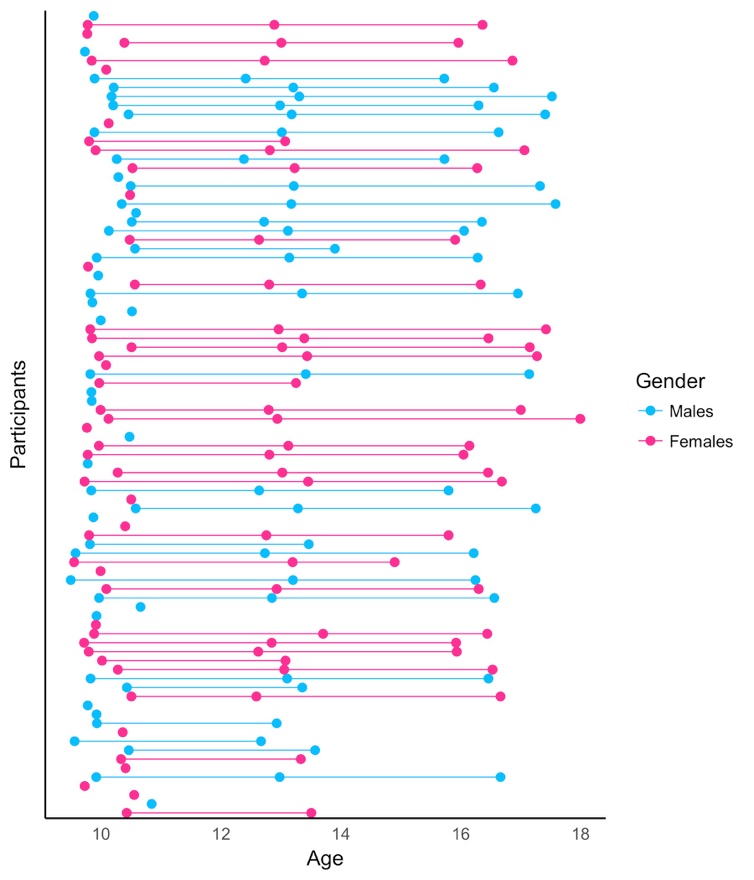

*Figure S1.* Representation of longitudinal sample.

Table S1. *SES,* *race & ethnicity*

|  | **Mean** | **SD** |
| --- | --- | --- |
| *SES* | 119 | 104 |
| *Race* | **N** | **%** |
| Caucasian | 62 | 69 |
| American Indian/ Alaskan Native | 5 | 6 |
| Pacific Islander / Asian | 5 | 6 |
| African American | 3 | 3 |
| Multiracial | 15 | 17 |
| *Hispanic Ethnicity* | 30 | 33 |

Note: SES was based on parent-reports of income. None of these demographic variables were related to PDS (SES: R = -0.148, p = 0.17; Race: F_(4/81)_ = 0.118, p = 0.97; Ethnicity: t_(55)_ = -1.76, p = 0.08)

Table S2. Attrition analyses – differences at baseline (wave1).

|  | Missing | Not missing |  |
| --- | --- | --- | --- |
| SES | 111.25 | 125.54 | t(88) = -0.65, p = 0.5 |
| Testosterone (log) | 2.98 | 3.16 | t(90) = -2, p = 0.1 |
| PDS | 1.83 | 1.84 | t(90) = -0.09, p = 0.9 |
| Age | 10.1 | 10 | t(90) = 0.9, p = 0.4 |
| Gender (M:F) | (24:20) | (21:50) | X^2^(1) = 0.4, p = 0.5 |

Note: Groups (missing vs not missing) are categorized based on inclusion in study at wave 3.

Table S3. Testosterone concentrations and intra-assay coefficient of variation (CV) at each wave.

|  |  |  | Concentrations | | | | Intra-assay CV | | | |
| --- | --- | --- | --- | --- | --- | --- | --- | --- | --- | --- |
|  | Wave | N | Mean | SD | Min | Max | Mean | SD | Min | Max |
| Males | 1 | 45 | 22.6 | 10.7 | 6.87 | 54.6 | 4.41 | 4.14 | 0 | 19.8 |
|  | 2 | 27 | 69.1 | 40.5 | 13.18 | 176.4 | 2.22 | 2.29 | 0.128 | 10.4 |
|  | 3 | 21 | 149.1 | 42.9 | 77.45 | 237.6 | 7.66 | 3.23 | 0.205 | 12 |
| Females | 1 | 45 | 25.8 | 11.9 | 5.25 | 57.1 | 4.09 | 7.19 | 0.115 | 47.8 |
|  | 2 | 30 | 54.8 | 49 | 12.63 | 275.7 | 3.03 | 2.51 | 0.187 | 11.7 |
|  | 3 | 25 | 68.6 | 32.2 | 32.25 | 178.6 | 5.97 | 3.08 | 0.612 | 11.3 |

Sex differences in (log) testosterone concentrations

Wave 1: t (90) = -1, p = 0.2

Wave 2: t (50) = 1, p = 0.1

Wave 3: t (40) - 8, p < 0.001

Table S4. ICC for hormones and ROIs across the three time points.

| Left Amygdala | 0.553 |
| --- | --- |
| Right Amygdala | 0.447 |
| Left Hippocampus | 0.502 |
| Right Hippocampus | 0.482 |
| Left NAcc | 0.405 |
| Right NAcc | 0.381 |
| Testosterone (females) | 0.30 |
| Testosterone (males) | 0.38 |

NB: Values reflect mean ICC for each of the five emotional conditions

**Table S5. BOLD response to the affective faces task at each wave.**

|  | Wave | Beta | SE | p |
| --- | --- | --- | --- | --- |
| Left Amygdala | 1 | 0.544 | 0.125 | <0.001 |
|  | 2 | 0.747 | 0.15 | <0.001 |
|  | 3 | 0.534 | 0.116 | <0.001 |
| Right Amygdala | 1 | 0.609 | 0.131 | <0.001 |
|  | 2 | 0.74 | 0.156 | <0.001 |
|  | 3 | 0.536 | 0.15 | <0.001 |
| Left Nacc | 1 | -0.185 | 0.084 | 0.028 |
|  | 2 | 0.046 | 0.129 | 0.723 |
|  | 3 | -0.191 | 0.139 | 0.172 |
| Right Nacc | 1 | -0.272 | 0.077 | <0.001 |
|  | 2 | -0.02 | 0.135 | 0.879 |
|  | 3 | -0.261 | 0.128 | 0.043 |
| Left Hippocampus | 1 | 0.26 | 0.084 | 0.002 |
|  | 2 | 0.526 | 0.112 | <0.001 |
|  | 3 | 0.441 | 0.096 | <0.001 |
| Right Hippocampus | 1 | 0.317 | 0.089 | <0.001 |
|  | 2 | 0.555 | 0.112 | <0.001 |
|  | 3 | 0.436 | 0.115 | <0.001 |

Note: Analyses were conducted across all five emotional conditions.

**Table S6. Model fit statistics for ROI models of age, PDS, and testosterone**

|  | | | | Model | | DF | AIC | logLik | Test | Chisq | ChiDF | Pval |
| --- | --- | --- | --- | --- | --- | --- | --- | --- | --- | --- | --- | --- |
| **Age** | | | |  | |  |  |  |  |  |  |  |
| *Left Amygdala* | | | |  | |  |  |  |  |  |  |  |
| Null | | | | 1 | | 3 | 2602.49 | -1298.24 |  |  |  |  |
| Linear | | | | 2 | | 4 | 2604.26 | -1298.13 | 1 vs 2 | 0.23 | 1 | 0.631 |
| Quadratic | | | | 3 | | 5 | 2605.88 | -1297.94 | 2 vs 3 | 0.38 | 1 | 0.537 |
| **Null + random age** | | | | **4** | | **5** | **2557.92** | **-1273.96** | **1 vs 4** | **48.57** | **2** | **0.000** |
| Null + random (age + cond) | | | | 5 | | 23 | 2571.26 | -1262.63 | 4 vs 5 | 22.66 | 18 | 0.204 |
| Null + sex + random age | | | | 6 | | 6 | 2559.41 | -1273.7 | 4 vs 6 | 0.51 | 1 | 0.474 |
| Null + cond + random age | | | | 7 | | 9 | 2560.06 | -1271.03 | 4 vs 7 | 5.86 | 4 | 0.210 |
| *Right Amygdala* | | | |  | |  |  |  |  |  |  |  |
| Null | | | | 1 | | 3 | 2638.92 | -1316.46 |  |  |  |  |
| Linear | | | | 2 | | 4 | 2640.08 | -1316.04 | 1 vs 2 | 0.84 | 1 | 0.359 |
| Quadratic | | | | 3 | | 5 | 2641.92 | -1315.96 | 2 vs 3 | 0.16 | 1 | 0.693 |
| Null + random age | | | | 4 | | 5 | 2571.1 | -1280.55 | 1 vs 4 | 71.82 | 2 | 0.000 |
| Null + random (age + cond) | | | | 5 | | 23 | 2601.52 | -1277.76 | 4 vs 5 | 5.58 | 18 | 0.998 |
| Null + sex + random age | | | | 6 | | 6 | 2572.73 | -1280.36 | 4 vs 6 | 0.37 | 1 | 0.542 |
| **Null + cond + random age** | | | | **7** | | **9** | **2568.57** | **-1275.28** | **4 vs 7** | **10.53** | **4** | **0.032** |
| *Left Nacc* | | | |  | |  |  |  |  |  |  |  |
| Null | | | | 1 | | 3 | 2321.87 | -1157.94 |  |  |  |  |
| Linear | | | | 2 | | 4 | 2323.87 | -1157.94 | 1 vs 2 | 0.00 | 1 | 0.984 |
| Quadratic | | | | 3 | | 5 | 2321.59 | -1155.79 | 2 vs 3 | 4.28 | 1 | 0.038 |
| Null + random age | | | | 4 | | 5 | 2285.94 | -1137.97 | 1 vs 4 | 39.94 | 2 | 0.000 |
| Null + random (age + cond) | | | | 5 | | 23 | 2312.13 | -1133.06 | 4 vs 5 | 9.81 | 18 | 0.938 |
| **Null + sex + random age** | | | | **6** | | **6** | **2283.29** | **-1135.65** | **4 vs 6** | **4.64** | **1** | **0.031** |
| Null + cond + random age | | | | 7 | | 9 | 2288.26 | -1135.13 | 4 vs 7 | 5.67 | 4 | 0.225 |
| *Right Nacc* | | | |  | |  |  |  |  |  |  |  |
| Null | | | | 1 | | 3 | 2234.13 | -1114.06 |  |  |  |  |
| Linear | | | | 2 | | 4 | 2235.72 | -1113.86 | 1 vs 2 | 0.41 | 1 | 0.520 |
| Quadratic | | | | 3 | | 5 | 2229.65 | -1109.82 | 2 vs 3 | 8.07 | 1 | 0.005 |
| Quadratic + random age | | | | 4 | | 7 | 2195.77 | -1090.88 | 3 vs 4 | 37.88 | 2 | 0.000 |
| Quadratic + random (age + cond) | | | | 5 | | 25 | 2216.49 | -1083.24 | 4 vs 5 | 15.28 | 18 | 0.643 |
| Quadratic + sex + random age | | | | 6 | | 8 | 2196.01 | -1090 | 4 vs 6 | 1.76 | 1 | 0.185 |
| **Quadratic*sex + random age** | | | | **7** | | **10** | **2186.51** | **-1083.25** | **6 vs 7** | **13.50** | **2** | **0.001** |
| *Left Hippocampus* | | | |  | |  |  |  |  |  |  |  |
| Null | | | | 1 | | 3 | 2161.76 | -1077.88 |  |  |  |  |
| Linear | | | | 2 | | 4 | 2160.29 | -1076.14 | 1 vs 2 | 3.47 | 1 | 0.063 |
| Quadratic | | | | 3 | | 5 | 2160.27 | -1075.14 | 2 vs 3 | 2.02 | 1 | 0.156 |
| Null + random age | | | | 4 | | 5 | 2126.89 | -1058.44 | 1 vs 4 | 38.87 | 2 | 0.000 |
| Null + random (age + cond) | | | | 5 | | 23 | 2141.6 | -1047.8 | 4 vs 5 | 21.29 | 18 | 0.265 |
| Null + sex + random age | | | | 6 | | 6 | 2128.03 | -1058.02 | 4 vs 6 | 0.85 | 1 | 0.356 |
| **Null + cond + random age** | | | | **7** | | **9** | **2124.72** | **-1053.36** | **4 vs 7** | **10.16** | **4** | **0.038** |
| *Right Hippocampus* | | | |  | |  |  |  |  |  |  |  |
| Null | | | | 1 | | 3 | 2159.43 | -1076.71 |  |  |  |  |
| Linear | | | | 2 | | 4 | 2160.85 | -1076.43 | 1 vs 2 | 0.58 | 1 | 0.448 |
| Quadratic | | | | 3 | | 5 | 2158.62 | -1074.31 | 2 vs 3 | 4.24 | 1 | 0.040 |
| **Null + random age** | | | | **4** | | **5** | **2100.92** | **-1045.46** | **3 vs 4** | **62.51** | **2** | **0.000** |
| Null + random (age + cond) | | | | 5 | | 23 | 2129.51 | -1041.76 | 4 vs 5 | 7.41 | 18 | 0.986 |
| Null + sex + random age | | | | 6 | | 6 | 2102.91 | -1045.46 | 4 vs 6 | 0.01 | 1 | 0.938 |
| Null + cond + random age | | | | 7 | | 9 | 2103.24 | -1042.62 | 4 vs 7 | 5.68 | 4 | 0.224 |
| **PDS** | | | |  | |  |  |  |  |  |  |  |
| *Left Amygdala* | | | |  | |  |  |  |  |  |  |  |
| Null | | | | 1 | | 3 | 2602.49 | -1298.24 |  |  |  |  |
| Linear | | | | 2 | | 4 | 2604.38 | -1298.19 | 1 vs 2 | 0.11 | 1 | 0.743 |
| Quadratic | | | | 3 | | 5 | 2605.53 | -1297.76 | 2 vs 3 | 0.85 | 1 | 0.356 |
| **Null + random age** | | | | **4** | | 5 | 2588.4 | -1289.2 | **1 vs 4** | 18.08 | 2 | 0.000 |
| Null + random (age + cond) | | | | 5 | | 23 | 2607.86 | -1280.93 | 4 vs 5 | 16.54 | 18 | 0.555 |
| Null + sex + random age | | | | 6 | | 6 | 2589.07 | -1288.53 | 4 vs 6 | 1.34 | 1 | 0.248 |
| Null + cond + random age | | | | 7 | | 9 | 2590.96 | -1286.48 | 4 vs 7 | 5.45 | 4 | 0.244 |
| *Right Amygdala* | | | |  | |  |  |  |  |  |  |  |
| Null | | | | 1 | | 3 | 2638.92 | -1316.46 |  |  |  |  |
| Linear | | | | 2 | | 4 | 2637.25 | -1314.62 | 1 vs 2 | 3.67 | 1 | 0.055 |
| Quadratic | | | | 3 | | 5 | 2637.13 | -1313.56 | 2 vs 3 | 2.12 | 1 | 0.145 |
| Null + random age | | | | 4 | | 5 | 2569.36 | -1279.68 | 1 vs 4 | 73.56 | 2 | 0.000 |
| Null + random (age + cond) | | | | 5 | | 23 | 2598.43 | -1276.22 | 4 vs 5 | 6.93 | 18 | 0.991 |
| Null + sex + random age | | | | 6 | | 6 | 2570.77 | -1279.39 | 4 vs 6 | 0.59 | 1 | 0.443 |
| **Null + cond + random age** | | | | **7** | | **9** | **2566.63** | **-1274.31** | **4 vs 7** | **10.73** | **4** | **0.030** |
| *Left Nacc* | | | |  | |  |  |  |  |  |  |  |
| Null | | | | 1 | | 3 | 2321.87 | -1157.94 |  |  |  |  |
| Linear | | | | 2 | | 4 | 2321.54 | -1156.77 | 1 vs 2 | 2.34 | 1 | 0.126 |
| Quadratic | | | | 3 | | 5 | 2323.03 | -1156.51 | 2 vs 3 | 0.51 | 1 | 0.476 |
| Null + random age | | | | **4** | | 5 | 2272.32 | -1131.16 | 1 vs 4 | 53.56 | 2 | 0.000 |
| Null + random (age + cond) | | | | 5 | | 23 | 2301.03 | -1127.51 | 4 vs 5 | 7.29 | 18 | 0.987 |
| **Null + sex + random age** | | | | **6** | | **6** | **2270.25** | **-1129.12** | **4 vs 6** | **4.07** | **1** | **0.044** |
| Null + cond + random age | | | | 7 | | 9 | 2274.51 | -1128.25 | 4 vs 7 | 5.81 | 4 | 0.214 |
| *Right Nacc* | | | |  | |  |  |  |  |  |  |  |
| Null | | | | 1 | | 3 | 2234.13 | -1114.06 |  |  |  |  |
| Linear | | | | 2 | | 4 | 2234.81 | -1113.4 | 1 vs 2 | 1.32 | 1 | 0.250 |
| Quadratic | | | | 3 | | 5 | 2236.64 | -1113.32 | 2 vs 3 | 0.16 | 1 | 0.686 |
| **Null + random age** | | | | **4** | | **5** | **2175.3** | **-1082.65** | **1 vs 4** | **62.83** | **2** | **0.000** |
| Null + random (age + cond) | | | | 5 | | 23 | 2202.45 | -1078.23 | 4 vs 5 | 8.85 | 18 | 0.963 |
| Null + sex + random age | | | | 6 | | 6 | 2176.18 | -1082.09 | 4 vs 6 | 1.13 | 1 | 0.289 |
| Null + cond + random age | | | | 7 | | 9 | 2178.38 | -1080.19 | 4 vs 7 | 4.93 | 4 | 0.295 |
| *Left Hippocampus* | | | |  | |  |  |  |  |  |  |  |
| Null | | | | 1 | | 3 | 2161.76 | -1077.88 |  |  |  |  |
| Linear | | | | 2 | | 4 | 2163.76 | -1077.88 | 1 vs 2 | 0.00 | 1 | 0.973 |
| Quadratic | | | | 3 | | 5 | 2165.39 | -1077.69 | 2 vs 3 | 0.37 | 1 | 0.543 |
| **Null + random age** | | | | **4** | | **5** | **2138.83** | **-1064.42** | **3 vs 4** | **26.92** | **2** | **0.000** |
| Null + random (age + cond) | | | | 5 | | 23 | 2159.21 | -1056.61 | 4 vs 5 | 15.62 | 18 | 0.619 |
| Null + sex + random age | | | | 6 | | 6 | 2139.91 | -1063.95 | 4 vs 6 | 0.93 | 1 | 0.335 |
| Null + cond + random age | | | | 7 | | 9 | 2137.04 | -1059.52 | 4 vs 7 | 9.79 | 4 | 0.044 |
| *Right Hippocampus* | | | |  | |  |  |  |  |  |  |  |
| Null | | | | 1 | | 3 | 2159.43 | -1076.71 |  |  |  |  |
| Linear | | | | 2 | | 4 | 2159.99 | -1075.99 | 1 vs 2 | 1.44 | 1 | 0.229 |
| Quadratic | | | | 3 | | 5 | 2159.91 | -1074.95 | 2 vs 3 | 2.08 | 1 | 0.149 |
| **Null + random age** | | | | **4** | | **5** | **2107.66** | **-1048.83** | **1 vs 4** | **55.77** | **2** | **0.000** |
| Null + random (age + cond) | | | | 5 | | 23 | 2136.34 | -1045.17 | 4 vs 5 | 7.32 | 18 | 0.987 |
| Null + sex + random age | | | | 6 | | 6 | 2109.63 | -1048.81 | 4 vs 6 | 0.03 | 1 | 0.862 |
| Null + cond + random age | | | | 7 | | 9 | 2110.11 | -1046.05 | 4 vs 7 | 5.55 | 4 | 0.236 |
| **Testosterone (females)** | | | |  | |  |  |  |  |  |  |  |
| *Left Amygdala* | | | |  | |  |  |  |  |  |  |  |
| Null | | | | 1 | | 3 | 1290.54 | -642.27 |  |  |  |  |
| Linear | | | | 2 | | 4 | 1292.53 | -642.26 | 1 vs 2 | 0.02 | 1 | 0.898 |
| Quadratic | | | | 3 | | 5 | 1285.14 | -637.57 | 2 vs 3 | 9.39 | 1 | 0.002 |
| **Quadratic + random age** | | | | **4** | | **7** | **1280.38** | **-633.19** | **3 vs 4** | **8.76** | **2** | **0.013** |
| Quadratic + random (age + cond) | | | | 5 | | 25 | 1302.63 | -626.32 | 4 vs 5 | 13.75 | 18 | 0.745 |
| Quadratic + cond + random age | | | | 6 | | 11 | 1280.08 | -629.04 | 4 vs 6 | 8.30 | 4 | 0.081 |
| Quadratic * cond + random age | | | | 7 | | 19 | 1288.97 | -625.49 | 6 vs 7 | 7.11 | 8 | 0.525 |
| *Right Amygdala* | | | |  | |  |  |  |  |  |  |  |
| Null | | | | 1 | | 3 | 1410.45 | -702.23 |  |  |  |  |
| Linear | | | | 2 | | 4 | 1412.28 | -702.14 | 1 vs 2 | 0.17 | 1 | 0.678 |
| Quadratic | | | | 3 | | 5 | 1408.43 | -699.22 | 2 vs 3 | 5.85 | 1 | 0.016 |
| **Quadratic + random age** | | | | **4** | | **7** | **1366.25** | **-676.12** | **3 vs 4** | **46.18** | **2** | **0.000** |
| Quadratic + random (age + cond) | | | | 5 | | 25 | 1394.7 | -672.35 | 4 vs 5 | 7.55 | 18 | 0.985 |
| Quadratic + cond + random age | | | | 6 | | 11 | 1371.79 | -674.9 | 4 vs 6 | 2.46 | 4 | 0.652 |
| Quadratic * cond + random age | | | | 7 | | 19 | 1381.23 | -671.62 | 6 vs 7 | 6.56 | 8 | 0.585 |
| *Left Nacc* | | | |  | |  |  |  |  |  |  |  |
| Null | | | | 1 | | 3 | 1094.68 | -544.34 |  |  |  |  |
| Linear | | | | 2 | | 4 | 1094.95 | -543.48 | 1 vs 2 | 1.73 | 1 | 0.189 |
| Quadratic | | | | 3 | | 5 | 1096.95 | -543.48 | 2 vs 3 | 0.00 | 1 | 0.996 |
| **Null + random age** | | | | **4** | | **5** | **1083.56** | **-536.78** | **1 vs 4** | **15.13** | **2** | **0.001** |
| Null + random (age + cond) | | | | 5 | | 23 | 1115.12 | -534.56 | 4 vs 5 | 4.44 | 18 | 0.999 |
| Null + cond + random age | | | | 6 | | 9 | 1089.62 | -535.81 | 4 vs 6 | 1.94 | 4 | 0.747 |
| *Right Nacc* | | | |  | |  |  |  |  |  |  |  |
| Null | | | | 1 | | 3 | 1020.29 | -507.15 |  |  |  |  |
| Linear | | | | 2 | | 4 | 1022.02 | -507.01 | 1 vs 2 | 0.28 | 1 | 0.600 |
| Quadratic | | | | 3 | | 5 | 1023.86 | -506.93 | 2 vs 3 | 0.16 | 1 | 0.690 |
| **Null + random age** | | | | **4** | | **5** | **1003.46** | **-496.73** | **1 vs 4** | **20.83** | **2** | **0.000** |
| Null + random (age + cond) | | | | 5 | | 23 | 1027.47 | -490.73 | 4 vs 5 | 11.99 | 18 | 0.848 |
| Null + cond + random age | | | | 6 | | 9 | 1009.93 | -495.97 | 4 vs 6 | 1.52 | 4 | 0.822 |
| *Left Hippocampus* | | | |  | |  |  |  |  |  |  |  |
| Null | | | | 1 | | 3 | 1046.36 | -520.18 |  |  |  |  |
| Linear | | | | 2 | | 4 | 1048.21 | -520.1 | 1 vs 2 | 0.15 | 1 | 0.694 |
| Quadratic | | | | 3 | | 5 | 1047.52 | -518.76 | 2 vs 3 | 2.69 | 1 | 0.101 |
| **Null + random age** | | | | **4** | | **5** | **1035.92** | **-512.96** | **1 vs 4** | **14.44** | **2** | **0.001** |
| Null + random (age + cond) | | | | 5 | | 23 | 1059.37 | -506.68 | 4 vs 5 | 12.55 | 18 | 0.818 |
| Null + cond + random age | | | | 6 | | 9 | 1034.55 | -508.27 | 4 vs 6 | 9.37 | 4 | 0.052 |
| *Right Hippocampus* | | | |  | |  |  |  |  |  |  |  |
| Null | | | | 1 | | 3 | 1068.63 | -531.32 |  |  |  |  |
| Linear | | | | 2 | | 4 | 1070.51 | -531.25 | 1 vs 2 | 0.13 | 1 | 0.721 |
| Quadratic | | | | 3 | | 5 | 1065.97 | -527.99 | 2 vs 3 | 6.53 | 1 | 0.011 |
| **Quadratic + random age** | | | | **4** | | **7** | **1034.83** | **-510.42** | **3 vs 4** | **35.14** | **2** | **0.000** |
| Quadratic + random (age + cond) | | | | 5 | | 25 | 1064.17 | -507.08 | 4 vs 5 | 6.67 | 18 | 0.993 |
| Quadratic + cond + random age | | | | 6 | | 11 | 1038.44 | -508.22 | 4 vs 6 | 4.39 | 4 | 0.356 |
| Quadratic * cond + random age | | | | 7 | | 19 | 1046.17 | -504.09 | 6 vs 7 | 8.27 | 8 | 0.407 |
| **Testosterone ( males)** | | | |  | |  |  |  |  |  |  |  |
| *Left Amygdala* | | | |  | |  |  |  |  |  |  |  |
| Null | | | | 1 | | 3 | 1310.23 | -652.11 |  |  |  |  |
| Linear | | | | 2 | | 4 | 1312.06 | -652.03 | 1 vs 2 | 0.17 | 1 | 0.680 |
| Quadratic | | | | 3 | | 5 | 1313.88 | -651.94 | 2 vs 3 | 0.17 | 1 | 0.677 |
| **Null + random age** | | | | **4** | | **5** | **1302.28** | **-646.14** | **1 vs 4** | **11.94** | **2** | **0.003** |
| Null + random (age + cond) | | | | 5 | | 23 | 1317.2 | -635.6 | 4 vs 5 | 21.09 | 18 | 0.275 |
| Null + cond + random age | | | | 6 | | 9 | 1308.11 | -645.05 | 4 vs 6 | 2.18 | 4 | 0.703 |
| *Right Amygdala* | | | |  | |  |  |  |  |  |  |  |
| Null | | | | 1 | | 3 | 1226.38 | -610.19 |  |  |  |  |
| Linear | | | | 2 | | 4 | 1228.09 | -610.05 | 1 vs 2 | 0.29 | 1 | 0.590 |
| Quadratic | | | | 3 | | 5 | 1226.36 | -608.18 | 2 vs 3 | 3.73 | 1 | 0.053 |
| Null + random age | | | | 4 | | 5 | 1219.06 | -604.53 | 1 vs 4 | 11.32 | 2 | 0.003 |
| Null + random (age + cond) | | | | 5 | | 23 | 1247.27 | -600.63 | 4 vs 5 | 7.79 | 18 | 0.982 |
| **Null + cond + random age** | | | | **6** | | **9** | **1215.01** | **-598.51** | **4 vs 6** | **12.05** | **4** | **0.017** |
| *Left Nacc* | | | |  | |  |  |  |  |  |  |  |
| Null | | | | 1 | | 3 | 1206.97 | -600.49 |  |  |  |  |
| Linear | | | | 2 | | 4 | 1208.49 | -600.25 | 1 vs 2 | 0.48 | 1 | 0.488 |
| Quadratic | | | | 3 | | 5 | 1210.33 | -600.17 | 2 vs 3 | 0.16 | 1 | 0.686 |
| **Null + random age** | | | | **4** | | **5** | **1188.09** | **-589.05** | **1 vs 4** | **22.88** | **2** | **0.000** |
| Null + random (age + cond) | | | | 5 | | 23 | 1210.95 | -582.47 | 4 vs 5 | 13.15 | 18 | 0.783 |
| Null + cond + random age | | | | 6 | | 9 | 1188.13 | -585.06 | 4 vs 6 | 7.97 | 4 | 0.093 |
| *Right Nacc* | | | |  | |  |  |  |  |  |  |  |
| Null | | | | 1 | | 3 | 1184.7 | -589.35 |  |  |  |  |
| Linear | | | | 2 | | 4 | 1185.8 | -588.9 | 1 vs 2 | 0.90 | 1 | 0.342 |
| Quadratic | | | | 3 | | 5 | 1187.79 | -588.89 | 2 vs 3 | 0.01 | 1 | 0.911 |
| **Null + random age** | | | | **4** | | **5** | **1163.43** | **-576.72** | **1 vs 4** | **25.27** | **2** | **0.000** |
| Null + random (age + cond) | | | | 5 | | 23 | 1186.66 | -570.33 | 4 vs 5 | 12.77 | 18 | 0.805 |
| Null + cond + random age | | | | 6 | | 9 | 1166.19 | -574.1 | 4 vs 6 | 5.24 | 4 | 0.264 |
| *Left Hippocampus* | | | |  | |  |  |  |  |  |  |  |
| Null | | | | 1 | | 3 | 1109.59 | -551.8 |  |  |  |  |
| Linear | | | | 2 | | 4 | 1110.21 | -551.1 | 1 vs 2 | 1.39 | 1 | 0.239 |
| Quadratic | | | | 3 | | 5 | 1110.87 | -550.44 | 2 vs 3 | 1.33 | 1 | 0.249 |
| **Null + random age** | | | | **4** | | **5** | **1104.12** | **-547.06** | **1 vs 4** | **9.48** | **2** | **0.009** |
| Null + random (age + cond) | | | | 5 | | 23 | 1121.92 | -537.96 | 4 vs 5 | 18.19 | 18 | 0.443 |
| Null + cond + random age | | | | 6 | | 9 | 1106.89 | -544.44 | 4 vs 6 | 5.23 | 4 | 0.265 |
| Null | | | | 1 | | 3 | 1109.59 | -551.8 |  |  |  |  |
| *Right Hippocampus* | | | |  | |  |  |  |  |  |  |  |
| Null | | | | 1 | | 3 | 1090.92 | -542.46 |  |  |  |  |
| Linear | | | | 2 | | 4 | 1092.86 | -542.43 | 1 vs 2 | 0.06 | 1 | 0.803 |
| Quadratic | | | | 3 | | 5 | 1093.98 | -541.99 | 2 vs 3 | 0.88 | 1 | 0.347 |
| **Null + random age** | | | | **4** | | **5** | **1075.58** | **-532.79** | **1 vs 4** | **19.34** | **2** | **0.000** |
| Null + random (age + cond) | | | | 5 | | 23 | 1105.53 | -529.76 | 4 vs 5 | 6.06 | 18 | 0.996 |
| Null + cond + random age | | | | 6 | | 9 | 1078.51 | -530.26 | 4 vs 6 | 5.07 | 4 | 0.280 |

**Table S7. Model fit statistics for ROI models of each emotional condition**

|  |  | **Model** | **DF** | **AIC** | **logLik** | **Test** | **Chisq** | | **ChiDF** | | **Pval** |
| --- | --- | --- | --- | --- | --- | --- | --- | --- | --- | --- | --- |
| **Age** | |  |  |  |  |  | |  | |  | |
| *Left Amgdala* | |  |  |  |  |  |  | |  | |  |
| Angry | **Null** | **1** | **3** | **552.71** | **-273.36** |  |  | |  | |  |
|  | Linear | 2 | 4 | 554.28 | -273.14 | 1 vs 2 | 0.43 | | 1 | | 0.511 |
|  | Quadratic | 3 | 5 | 556.25 | -273.12 | 2 vs 3 | 0.03 | | 1 | | 0.855 |
|  | Null + Sex | 4 | 4 | 554.31 | -273.15 | 1 vs 4 | 0.41 | | 1 | | 0.524 |
| Fear | **Null** | **1** | **3** | **571.99** | **-282.99** |  |  | |  | |  |
|  | Linear | 2 | 4 | 573.8 | -282.9 | 1 vs 2 | 0.18 | | 1 | | 0.667 |
|  | Quadratic | 3 | 5 | 575.4 | -282.7 | 2 vs 3 | 0.4 | | 1 | | 0.526 |
|  | Null + Sex | 4 | 4 | 571.87 | -281.94 | 1 vs 4 | 2.12 | | 1 | | 0.146 |
| Sad | **Null** | **1** | **3** | **558.66** | **-276.33** |  |  | |  | |  |
|  | Linear | 2 | 4 | 560.58 | -276.29 | 1 vs 2 | 0.09 | | 1 | | 0.767 |
|  | Quadratic | 3 | 5 | 558.24 | -274.12 | 2 vs 3 | 4.34 | | 1 | | 0.037 |
|  | Null + Sex | 4 | 4 | 560.23 | -276.11 | 1 vs 4 | 0.44 | | 1 | | 0.508 |
| Happy | **Null** | **1** | **3** | **531.85** | **-262.92** |  |  | |  | |  |
|  | Linear | 2 | 4 | 533.83 | -262.92 | 1 vs 2 | 0.01 | | 1 | | 0.908 |
|  | Quadratic | 3 | 5 | 534.77 | -262.39 | 2 vs 3 | 1.06 | | 1 | | 0.304 |
|  | Null + Sex | 4 | 4 | 533.84 | -262.92 | 1 vs 4 | 0.01 | | 1 | | 0.942 |
| Neutral | **Null** | **1** | **3** | **516.11** | **-255.06** |  |  | |  | |  |
|  | Linear | 2 | 4 | 517.22 | -254.61 | 1 vs 2 | 0.89 | | 1 | | 0.346 |
|  | Quadratic | 3 | 5 | 519.22 | -254.61 | 2 vs 3 | 0.01 | | 1 | | 0.942 |
|  | Null + Sex | 4 | 4 | 515.07 | -253.54 | 1 vs 4 | 3.04 | | 1 | | 0.081 |
| *Right Amygdala* | |  |  |  |  |  |  | |  | |  |
| Angry | **Null** | **1** | **3** | **575.03** | **-284.51** |  |  | |  | |  |
|  | Linear | 2 | 4 | 577.01 | -284.51 | 1 vs 2 | 0.01 | | 1 | | 0.917 |
|  | Quadratic | 3 | 5 | 578.96 | -284.48 | 2 vs 3 | 0.05 | | 1 | | 0.818 |
|  | Null + Sex | 4 | 4 | 574.53 | -283.27 | 1 vs 4 | 2.49 | | 1 | | 0.114 |
| Fear | **Null** | **1** | **3** | **554.26** | **-274.13** |  |  | |  | |  |
|  | Linear | 2 | 4 | 556.19 | -274.1 | 1 vs 2 | 0.07 | | 1 | | 0.798 |
|  | Quadratic | 3 | 5 | 557.82 | -273.91 | 2 vs 3 | 0.37 | | 1 | | 0.544 |
|  | Null + Sex | 4 | 4 | 554.38 | -273.19 | 1 vs 4 | 1.88 | | 1 | | 0.171 |
| Sad | **Null** | **1** | **3** | **543.13** | **-268.56** |  |  | |  | |  |
|  | Linear | 2 | 4 | 545.09 | -268.55 | 1 vs 2 | 0.04 | | 1 | | 0.851 |
|  | Quadratic | 3 | 5 | 546.23 | -268.12 | 2 vs 3 | 0.86 | | 1 | | 0.354 |
|  | Null + Sex | 4 | 4 | 544.72 | -268.36 | 1 vs 4 | 0.41 | | 1 | | 0.524 |
| Happy | **Null** | **1** | **3** | **547.68** | **-270.84** |  |  | |  | |  |
|  | Linear | 2 | 4 | 548.65 | -270.32 | 1 vs 2 | 1.03 | | 1 | | 0.31 |
|  | Quadratic | 3 | 5 | 549.46 | -269.73 | 2 vs 3 | 1.18 | | 1 | | 0.277 |
|  | Null + Sex | 4 | 4 | 549.67 | -270.84 | 1 vs 4 | 0 | | 1 | | 0.961 |
| Neutral | **Null** | **1** | **3** | **557.3** | **-275.65** |  |  | |  | |  |
|  | Linear | 2 | 4 | 558.82 | -275.41 | 1 vs 2 | 0.48 | | 1 | | 0.49 |
|  | Quadratic | 3 | 5 | 560.81 | -275.41 | 2 vs 3 | 0.01 | | 1 | | 0.912 |
|  | Null + Sex | 4 | 4 | 557.75 | -274.87 | 1 vs 4 | 1.55 | | 1 | | 0.213 |
| *Left NAcc* |  |  |  |  |  |  |  | |  | |  |
| Angry | Null | 1 | 3 | 484.7 | -239.35 |  |  | |  | |  |
|  | Linear | 2 | 4 | 485.53 | -238.76 | 1 vs 2 | 1.17 | | 1 | | 0.279 |
|  | Quadratic | 3 | 5 | 487.25 | -238.62 | 2 vs 3 | 0.28 | | 1 | | 0.597 |
|  | **Null + Sex** | **4** | **4** | **478.74** | **-235.37** | **1 vs 4** | **7.96** | | **1** | | **0.005** |
| Fear | Null | 1 | 3 | 475.29 | -234.65 |  |  | |  | |  |
|  | Linear | 2 | 4 | 477.1 | -234.55 | 1 vs 2 | 0.19 | | 1 | | 0.664 |
|  | **Quadratic** | **3** | **5** | **472.29** | **-231.15** | **2 vs 3** | **6.81** | | **1** | | **0.009** |
|  | Quadratic + Sex | 4 | 6 | 473.54 | -230.77 | 3 vs 4 | 0.75 | | 1 | | 0.387 |
|  | Quadratic*Sex | 5 | 8 | 477.46 | -230.73 | 4 vs 5 | 0.09 | | 2 | | 0.958 |
| Sad | Null | 1 | 3 | 496.01 | -245 |  |  | |  | |  |
|  | Linear | 2 | 4 | 496.36 | -244.18 | 1 vs 2 | 1.65 | | 1 | | 0.199 |
|  | Quadratic | 3 | 5 | 497.84 | -243.92 | 2 vs 3 | 0.52 | | 1 | | 0.472 |
|  | **Null + Sex** | **4** | **4** | **492.61** | **-242.3** | **1 vs 4** | **5.4** | | **1** | | **0.02** |
| Happy | **Null** | **1** | **3** | **470.26** | **-232.13** |  |  | |  | |  |
|  | Linear | 2 | 4 | 471.67 | -231.83 | 1 vs 2 | 0.59 | | 1 | | 0.442 |
|  | Quadratic | 3 | 5 | 472.65 | -231.32 | 2 vs 3 | 1.02 | | 1 | | 0.312 |
|  | Null + Sex | 4 | 4 | 471.33 | -231.66 | 1 vs 4 | 0.93 | | 1 | | 0.334 |
| Neutral | Null | 1 | 3 | 506.2 | -250.1 |  |  | |  | |  |
|  | Linear | 2 | 4 | 508.2 | -250.1 | 1 vs 2 | 0 | | 1 | | 0.967 |
|  | Quadratic | 3 | 5 | 509.56 | -249.78 | 2 vs 3 | 0.64 | | 1 | | 0.425 |
|  | **Null + Sex** | **4** | **4** | **503.75** | **-247.88** | **1 vs 4** | **4.45** | | **1** | | **0.035** |
| *Right NAcc* |  |  |  |  |  |  |  | |  | |  |
| Angry | Null | 1 | 3 | 466.59 | -230.3 |  |  | |  | |  |
|  | Linear | 2 | 4 | 467.41 | -229.7 | 1 vs 2 | 1.19 | | 1 | | 0.276 |
|  | Quadratic | 3 | 5 | 468.48 | -229.24 | 2 vs 3 | 0.93 | | 1 | | 0.336 |
|  | **Null + Sex** | **4** | **4** | **463.25** | **-227.63** | **1 vs 4** | **5.34** | | **1** | | **0.021** |
| Fear | **Null** | **1** | **3** | **443.69** | **-218.84** |  |  | |  | |  |
|  | Linear | 2 | 4 | 445.62 | -218.81 | 1 vs 2 | 0.06 | | 1 | | 0.801 |
|  | Quadratic | 3 | 5 | 444.86 | -217.43 | 2 vs 3 | 2.76 | | 1 | | 0.096 |
|  | Null + Sex | 4 | 4 | 445.24 | -218.62 | 3 vs 4 | 0.45 | | 1 | | 0.503 |
| Sad | **Null** | **1** | **3** | **497.68** | **-245.84** |  |  | |  | |  |
|  | Linear | 2 | 4 | 497.71 | -244.85 | 1 vs 2 | 1.98 | | 1 | | 0.16 |
|  | Quadratic | 3 | 5 | 496.82 | -243.41 | 2 vs 3 | 2.89 | | 1 | | 0.089 |
|  | Null + Sex | 4 | 4 | 498.89 | -245.45 | 1 vs 4 | 0.79 | | 1 | | 0.375 |
| Happy | **Null** | **1** | **3** | **445.43** | **-219.71** |  |  | |  | |  |
|  | Linear | 2 | 4 | 447.16 | -219.58 | 1 vs 2 | 0.27 | | 1 | | 0.604 |
|  | Quadratic | 3 | 5 | 447.16 | -218.58 | 2 vs 3 | 2 | | 1 | | 0.157 |
|  | Null + Sex | 4 | 4 | 446.47 | -219.24 | 1 vs 4 | 0.96 | | 1 | | 0.328 |
| Neutral | **Null** | **1** | **3** | **478.85** | **-236.42** |  |  | |  | |  |
|  | Linear | 2 | 4 | 480.6 | -236.3 | 1 vs 2 | 0.24 | | 1 | | 0.621 |
|  | Quadratic | 3 | 5 | 482.33 | -236.16 | 2 vs 3 | 0.28 | | 1 | | 0.599 |
|  | Null + Sex | 4 | 4 | 479.24 | -235.62 | 1 vs 4 | 1.6 | | 1 | | 0.205 |
| *Left Hippocampus* | |  |  |  |  |  |  | |  | |  |
| Angry | Null | 1 | 3 | 440.88 | -217.44 |  |  | |  | |  |
|  | Linear | 2 | 4 | 442.84 | -217.42 | 1 vs 2 | 0.04 | | 1 | | 0.84 |
|  | Quadratic | 3 | 5 | 444.79 | -217.4 | 2 vs 3 | 0.05 | | 1 | | 0.827 |
|  | Null + Sex | 4 | 4 | 442.36 | -217.18 | 1 vs 4 | 0.52 | | 1 | | 0.47 |
| Fear | **Null** | **1** | **3** | **481.17** | **-237.59** |  |  | |  | |  |
|  | Linear | 2 | 4 | 481.31 | -236.65 | 1 vs 2 | 1.87 | | 1 | | 0.172 |
|  | Quadratic | 3 | 5 | 482.85 | -236.43 | 2 vs 3 | 0.45 | | 1 | | 0.501 |
|  | Null + Sex | 4 | 4 | 480.83 | -236.41 | 1 vs 4 | 2.34 | | 1 | | 0.126 |
| Sad | Null | 1 | 3 | 445.9 | -219.95 |  |  | |  | |  |
|  | Linear | 2 | 4 | 445.85 | -218.93 | 1 vs 2 | 2.04 | | 1 | | 0.153 |
|  | **Quadratic** | **3** | **5** | **442.71** | **-216.36** | **2 vs 3** | **5.14** | | **1** | | **0.023** |
|  | Quadratic + Sex | 4 | 6 | 443.41 | -215.71 | 3 vs 4 | 1.3 | | 1 | | 0.254 |
|  | Quadratic*Sex | 5 | 8 | 446.97 | -215.48 | 4 vs 5 | 0.44 | | 2 | | 0.802 |
| Happy | **Null** | **1** | **3** | **446.25** | **-220.13** |  |  | |  | |  |
|  | Linear | 2 | 4 | 447.22 | -219.61 | 1 vs 2 | 1.03 | | 1 | | 0.31 |
|  | Quadratic | 3 | 5 | 448.68 | -219.34 | 2 vs 3 | 0.54 | | 1 | | 0.464 |
|  | Null + Sex | 4 | 4 | 448.14 | -220.07 | 1 vs 4 | 0.11 | | 1 | | 0.742 |
| Neutral | **Null** | **1** | **3** | **442.16** | **-218.08** |  |  | |  | |  |
|  | Linear | 2 | 4 | 441.96 | -216.98 | 1 vs 2 | 2.21 | | 1 | | 0.137 |
|  | Quadratic | 3 | 5 | 443.45 | -216.72 | 2 vs 3 | 0.51 | | 1 | | 0.475 |
|  | Null + Sex | 4 | 4 | 441.94 | -216.97 | 1 vs 4 | 2.22 | | 1 | | 0.136 |
| *Right Hippocampus* | | |  |  |  |  |  | |  | |  |
| Angry | **Null** | **1** | **3** | **484.87** | **-239.43** |  |  | |  | |  |
|  | Linear | 2 | 4 | 486.54 | -239.27 | 1 vs 2 | 0.32 | | 1 | | 0.569 |
|  | Quadratic | 3 | 5 | 488.31 | -239.15 | 2 vs 3 | 0.23 | | 1 | | 0.629 |
|  | Null + Sex | 4 | 4 | 486.45 | -239.23 | 1 vs 4 | 0.41 | | 1 | | 0.521 |
| Fear | **Null** | **1** | **3** | **454.92** | **-224.46** |  |  | |  | |  |
|  | Linear | 2 | 4 | 455.7 | -223.85 | 1 vs 2 | 1.22 | | 1 | | 0.27 |
|  | Quadratic | 3 | 5 | 457.44 | -223.72 | 2 vs 3 | 0.26 | | 1 | | 0.613 |
|  | Null + Sex | 4 | 4 | 455.83 | -223.92 | 3 vs 4 | 1.08 | | 1 | | 0.298 |
| Sad | **Null** | **1** | **3** | **431.75** | **-212.88** |  |  | |  | |  |
|  | Linear | 2 | 4 | 433.04 | -212.52 | 1 vs 2 | 0.71 | | 1 | | 0.398 |
|  | Quadratic | 3 | 5 | 431.13 | -210.56 | 2 vs 3 | 3.91 | | 1 | | 0.048 |
|  | Null + Sex | 4 | 4 | 433.4 | -212.7 | 1 vs 4 | 0.35 | | 1 | | 0.554 |
| Happy | **Null** | **1** | **3** | **425.95** | **-209.98** |  |  | |  | |  |
|  | Linear | 2 | 4 | 427.33 | -209.66 | 1 vs 2 | 0.63 | | 1 | | 0.429 |
|  | Quadratic | 3 | 5 | 427.16 | -208.58 | 2 vs 3 | 2.17 | | 1 | | 0.141 |
|  | Null + Sex | 4 | 4 | 426.93 | -209.47 | 1 vs 4 | 1.02 | | 1 | | 0.313 |
| Neutral | **Null** | **1** | **3** | **470.08** | **-232.04** |  |  | |  | |  |
|  | Linear | 2 | 4 | 472.04 | -232.02 | 1 vs 2 | 0.04 | | 1 | | 0.849 |
|  | Quadratic | 3 | 5 | 472.9 | -231.45 | 2 vs 3 | 1.15 | | 1 | | 0.285 |
|  | Null + Sex | 4 | 4 | 472.07 | -232.03 | 1 vs 4 | 0.01 | | 1 | | 0.925 |
| **PDS** | |  |  |  |  |  |  | |  | |  |
| *Left Amygdala* | |  |  |  |  |  |  | |  | |  |
| Angry | **Null** | **1** | **3** | **552.71** | **-273.36** |  |  | |  | |  |
|  | Linear | 2 | 4 | 552.58 | -272.29 | 1 vs 2 | 2.13 | | 1 | | 0.145 |
|  | Quadratic | 3 | 5 | 554.42 | -272.21 | 2 vs 3 | 0.16 | | 1 | | 0.686 |
|  | Null + Sex | 4 | 4 | 554.31 | -273.15 | 1 vs 4 | 0.41 | | 1 | | 0.524 |
| Fear | Null | 1 | 3 | 571.99 | -282.99 |  |  | |  | |  |
|  | Linear | 2 | 4 | 573.99 | -282.99 | 1 vs 2 | 0 | | 1 | | 0.978 |
|  | **Quadratic** | **3** | **5** | **564.77** | **-277.38** | **2 vs 3** | **11.22** | | **1** | | **0.001** |
|  | Quadratic + Sex | 4 | 6 | 565.86 | -276.93 | 3 vs 4 | 0.9 | | 1 | | 0.342 |
|  | Quadratic*Sex | 5 | 8 | 569.51 | -276.76 | 4 vs 5 | 0.35 | | 2 | | 0.838 |
| Sad | **Null** | **1** | **3** | **558.66** | **-276.33** |  |  | |  | |  |
|  | Linear | 2 | 4 | 560.62 | -276.31 | 1 vs 2 | 0.05 | | 1 | | 0.827 |
|  | Quadratic | 3 | 5 | 561.85 | -275.92 | 2 vs 3 | 0.77 | | 1 | | 0.38 |
|  | Null + Sex | 4 | 4 | 560.23 | -276.11 | 1 vs 4 | 0.44 | | 1 | | 0.508 |
| Happy | **Null** | **1** | **3** | **531.85** | **-262.92** |  |  | |  | |  |
|  | Linear | 2 | 4 | 533.59 | -262.79 | 1 vs 2 | 0.26 | | 1 | | 0.61 |
|  | Quadratic | 3 | 5 | 534.95 | -262.48 | 2 vs 3 | 0.63 | | 1 | | 0.427 |
|  | Null + Sex | 4 | 4 | 533.84 | -262.92 | 1 vs 4 | 0.01 | | 1 | | 0.942 |
| Neutral | **Null** | **1** | **3** | **516.11** | **-255.06** |  |  | |  | |  |
|  | Linear | 2 | 4 | 518.02 | -255.01 | 1 vs 2 | 0.09 | | 1 | | 0.759 |
|  | Quadratic | 3 | 5 | 520 | -255 | 2 vs 3 | 0.01 | | 1 | | 0.904 |
|  | Null + Sex | 4 | 4 | 515.07 | -253.54 | 1 vs 4 | 3.04 | | 1 | | 0.081 |
| *Right Amygdala* | |  |  |  |  |  |  | |  | |  |
| Angry | **Null** | **1** | **3** | **575.03** | **-284.51** |  |  | |  | |  |
|  | Linear | 2 | 4 | 575.34 | -283.67 | 1 vs 2 | 1.68 | | 1 | | 0.194 |
|  | Quadratic | 3 | 5 | 577.14 | -283.57 | 2 vs 3 | 0.2 | | 1 | | 0.653 |
|  | Null + Sex | 4 | 4 | 574.53 | -283.27 | 1 vs 4 | 2.49 | | 1 | | 0.114 |
| Fear | **Null** | **1** | **3** | **554.26** | **-274.13** |  |  | |  | |  |
|  | Linear | 2 | 4 | 556.17 | -274.08 | 1 vs 2 | 0.09 | | 1 | | 0.761 |
|  | Quadratic | 3 | 5 | 552.92 | -271.46 | 2 vs 3 | 5.24 | | 1 | | 0.022 |
|  | Null + Sex | 4 | 4 | 554.38 | -273.19 | 1 vs 4 | 1.88 | | 1 | | 0.171 |
| Sad | **Null** | **1** | **3** | **543.13** | **-268.56** |  |  | |  | |  |
|  | Linear | 2 | 4 | 545.06 | -268.53 | 1 vs 2 | 0.06 | | 1 | | 0.799 |
|  | Quadratic | 3 | 5 | 545.61 | -267.8 | 2 vs 3 | 1.46 | | 1 | | 0.228 |
|  | Null + Sex | 4 | 4 | 544.72 | -268.36 | 1 vs 4 | 0.41 | | 1 | | 0.524 |
| Happy | **Null** | **1** | **3** | **547.68** | **-270.84** |  |  | |  | |  |
|  | Linear | 2 | 4 | 548.47 | -270.23 | 1 vs 2 | 1.21 | | 1 | | 0.271 |
|  | Quadratic | 3 | 5 | 550.47 | -270.23 | 2 vs 3 | 0 | | 1 | | 0.965 |
|  | Null + Sex | 4 | 4 | 549.67 | -270.84 | 1 vs 4 | 0 | | 1 | | 0.961 |
| Neutral | **Null** | **1** | **3** | **557.3** | **-275.65** |  |  | |  | |  |
|  | Linear | 2 | 4 | 557.96 | -274.98 | 1 vs 2 | 1.34 | | 1 | | 0.247 |
|  | Quadratic | 3 | 5 | 558.5 | -274.25 | 2 vs 3 | 1.46 | | 1 | | 0.228 |
|  | Null + Sex | 4 | 4 | 557.75 | -274.87 | 1 vs 4 | 1.55 | | 1 | | 0.213 |
| *Left NAcc* |  |  |  |  |  |  |  | |  | |  |
| Angry | Null | 1 | 3 | 484.7 | -239.35 |  |  | |  | |  |
|  | Linear | 2 | 4 | 480.9 | -236.45 | 1 vs 2 | 5.8 | | 1 | | 0.016 |
|  | Quadratic | 3 | 5 | 482.57 | -236.29 | 2 vs 3 | 0.33 | | 1 | | 0.568 |
|  | **Linear + Sex** | 4 | **5** | **476.59** | **-233.29** | **2 vs 4** | **6.31** | | **1** | | **0.012** |
|  | Linear*Sex | **5** | 6 | 478.58 | -233.29 | 4 vs 5 | 0.01 | | 1 | | 0.938 |
| Fear | **Null** | **1** | **3** | **475.29** | **-234.65** |  |  | |  | |  |
|  | Linear | 2 | 4 | 477.28 | -234.64 | 1 vs 2 | 0.01 | | 1 | | 0.918 |
|  | Quadratic | 3 | 5 | 474.83 | -232.41 | 2 vs 3 | 4.45 | | 1 | | 0.035 |
|  | Null + Sex | 4 | 4 | 476.63 | -234.31 | 1 vs 4 | 0.67 | | 1 | | 0.415 |
| Sad | Null | 1 | 3 | 496.01 | -245 |  |  | |  | |  |
|  | Linear | 2 | 4 | 498.01 | -245 | 1 vs 2 | 0 | | 1 | | 0.998 |
|  | Quadratic | 3 | 5 | 498.66 | -244.33 | 2 vs 3 | 1.35 | | 1 | | 0.245 |
|  | **Null + Sex** | 4 | **4** | **492.61** | **-242.3** | **1 vs 4** | **5.4** | | **1** | | **0.02** |
| Happy | **Null** | **1** | **3** | **470.26** | **-232.13** |  |  | |  | |  |
|  | Linear | 2 | 4 | 472.25 | -232.13 | 1 vs 2 | 0.01 | | 1 | | 0.938 |
|  | Quadratic | 3 | 5 | 469.58 | -229.79 | 2 vs 3 | 4.67 | | 1 | | 0.031 |
|  | Null + Sex | 4 | 4 | 471.33 | -231.66 | 1 vs 4 | 0.93 | | 1 | | 0.334 |
| Neutral | Null | 1 | 3 | 506.2 | -250.1 |  |  | |  | |  |
|  | Linear | 2 | 4 | 507.51 | -249.75 | 1 vs 2 | 0.69 | | 1 | | 0.407 |
|  | Quadratic | 3 | 5 | 509.4 | -249.7 | 2 vs 3 | 0.11 | | 1 | | 0.736 |
|  | **Null + Sex** | 4 | **4** | **503.75** | **-247.88** | **1 vs 4** | **4.45** | | **1** | | **0.035** |
| *Right NAcc* |  |  |  |  |  |  |  | |  | |  |
| Angry | Null | 1 | 3 | 466.59 | -230.3 |  |  | |  | |  |
|  | **Linear** | **2** | **4** | **462.39** | **-227.19** | **1 vs 2** | **6.2** | | **1** | | **0.013** |
|  | Quadratic | 3 | 5 | 464.39 | -227.19 | 2 vs 3 | 0 | | 1 | | 0.996 |
|  | Linear + Sex | 4 | 5 | 460.41 | -225.21 | 2 vs 4 | 3.97 | | 1 | | 0.046 |
|  | Linear*Sex | 5 | 6 | 462.13 | -225.06 | 4 vs 5 | 0.29 | | 1 | | 0.592 |
| Fear | **Null** | **1** | **3** | **443.69** | **-218.84** |  |  | |  | |  |
|  | Linear | 2 | 4 | 445.45 | -218.73 | 1 vs 2 | 0.23 | | 1 | | 0.63 |
|  | Quadratic | 3 | 5 | 442.97 | -216.49 | 2 vs 3 | 4.48 | | 1 | | 0.034 |
|  | Null + Sex | 4 | 4 | 445.24 | -218.62 | 1 vs 4 | 0.45 | | 1 | | 0.503 |
| Sad | **Null** | **1** | **3** | **497.68** | **-245.84** |  |  | |  | |  |
|  | Linear | 2 | 4 | 499.2 | -245.6 | 1 vs 2 | 0.48 | | 1 | | 0.489 |
|  | Quadratic | 3 | 5 | 501.1 | -245.55 | 2 vs 3 | 0.1 | | 1 | | 0.752 |
|  | Null + Sex | 4 | 4 | 498.89 | -245.45 | 1 vs 4 | 0.79 | | 1 | | 0.375 |
| Happy | **Null** | **1** | **3** | **445.43** | **-219.71** |  |  | |  | |  |
|  | Linear | 2 | 4 | 446.79 | -219.4 | 1 vs 2 | 0.63 | | 1 | | 0.426 |
|  | Quadratic | 3 | 5 | 447.1 | -218.55 | 2 vs 3 | 1.69 | | 1 | | 0.193 |
|  | Null + Sex | 4 | 4 | 446.47 | -219.24 | 1 vs 4 | 0.96 | | 1 | | 0.328 |
| Neutral | **Null** | **1** | **3** | **478.85** | **-236.42** |  |  | |  | |  |
|  | Linear | 2 | 4 | 480.38 | -236.19 | 1 vs 2 | 0.47 | | 1 | | 0.495 |
|  | Quadratic | 3 | 5 | 482.15 | -236.07 | 2 vs 3 | 0.23 | | 1 | | 0.63 |
|  | Null + Sex | 4 | 4 | 479.24 | -235.62 | 1 vs 4 | 1.6 | | 1 | | 0.205 |
| *Left Hippocampus* | |  |  |  |  |  |  | |  | |  |
| Angry | **Null** | **1** | **3** | **440.88** | **-217.44** |  |  | |  | |  |
|  | Linear | 2 | 4 | 441.47 | -216.74 | 1 vs 2 | 1.41 | | 1 | | 0.235 |
|  | Quadratic | 3 | 5 | 443.44 | -216.72 | 2 vs 3 | 0.04 | | 1 | | 0.848 |
|  | Null + Sex | 4 | 4 | 442.36 | -217.18 | 1 vs 4 | 0.52 | | 1 | | 0.47 |
| Fear | Null | 1 | 3 | 481.17 | -237.59 |  |  | |  | |  |
|  | Linear | 2 | 4 | 482.86 | -237.43 | 1 vs 2 | 0.31 | | 1 | | 0.577 |
|  | **Quadratic** | **3** | **5** | **476.43** | **-233.21** | **2 vs 3** | **8.43** | | **1** | | **0.004** |
|  | Quadratic + Sex | 4 | 6 | 477.11 | -232.55 | 3 vs 4 | 1.32 | | 1 | | 0.25 |
|  | Quadratic*Sex | 5 | 8 | 479.9 | -231.95 | 4 vs 5 | 1.21 | | 2 | | 0.546 |
| Sad | **Null** | **1** | **3** | **445.9** | **-219.95** |  |  | |  | |  |
|  | Linear | 2 | 4 | 447.59 | -219.8 | 1 vs 2 | 0.31 | | 1 | | 0.58 |
|  | Quadratic | 3 | 5 | 449.59 | -219.8 | 2 vs 3 | 0 | | 1 | | 0.965 |
|  | Null + Sex | 4 | 4 | 446.71 | -219.36 | 1 vs 4 | 1.18 | | 1 | | 0.277 |
| Happy | **Null** | **1** | **3** | **446.25** | **-220.13** |  |  | |  | |  |
|  | Linear | 2 | 4 | 448.24 | -220.12 | 1 vs 2 | 0.01 | | 1 | | 0.934 |
|  | Quadratic | 3 | 5 | 449.94 | -219.97 | 2 vs 3 | 0.31 | | 1 | | 0.578 |
|  | Null + Sex | 4 | 4 | 448.14 | -220.07 | 1 vs 4 | 0.11 | | 1 | | 0.742 |
| Neutral | **Null** | **1** | **3** | **442.16** | **-218.08** |  |  | |  | |  |
|  | Linear | 2 | 4 | 444.15 | -218.07 | 1 vs 2 | 0.01 | | 1 | | 0.904 |
|  | Quadratic | 3 | 5 | 446.09 | -218.04 | 2 vs 3 | 0.06 | | 1 | | 0.806 |
|  | Null + Sex | 4 | 4 | 441.94 | -216.97 | 1 vs 4 | 2.22 | | 1 | | 0.136 |
| *Right Hippocampus* | |  |  |  |  |  |  | |  | |  |
| Angry | **Null** | **1** | **3** | **484.87** | **-239.43** |  |  | |  | |  |
|  | Linear | 2 | 4 | 486.15 | -239.07 | 1 vs 2 | 0.72 | | 1 | | 0.397 |
|  | Quadratic | 3 | 5 | 488.14 | -239.07 | 2 vs 3 | 0.01 | | 1 | | 0.94 |
|  | Null + Sex | 4 | 4 | 486.45 | -239.23 | 1 vs 4 | 0.41 | | 1 | | 0.521 |
| Fear | Null | 1 | 3 | 454.92 | -224.46 |  |  | |  | |  |
|  | Linear | 2 | 4 | 456.78 | -224.39 | 1 vs 2 | 0.13 | | 1 | | 0.717 |
|  | **Quadratic** | **3** | **5** | **449.06** | **-219.53** | **2 vs 3** | **9.73** | | **1** | | **0.002** |
|  | Quadratic + Sex | 4 | 6 | 450.73 | -219.37 | 3 vs 4 | 0.33 | | 1 | | 0.568 |
|  | Quadratic*Sex | 5 | 8 | 454.57 | -219.29 | 4 vs 5 | 0.16 | | 2 | | 0.923 |
| Sad | **Null** | **1** | **3** | **431.75** | **-212.88** |  |  | |  | |  |
|  | Linear | 2 | 4 | 433.66 | -212.83 | 1 vs 2 | 0.09 | | 1 | | 0.763 |
|  | Quadratic | 3 | 5 | 435.66 | -212.83 | 2 vs 3 | 0 | | 1 | | 0.989 |
|  | Null + Sex | 4 | 4 | 433.4 | -212.7 | 1 vs 4 | 0.35 | | 1 | | 0.554 |
| Happy | **Null** | **1** | **3** | **425.95** | **-209.98** |  |  | |  | |  |
|  | Linear | 2 | 4 | 427.71 | -209.86 | 1 vs 2 | 0.24 | | 1 | | 0.624 |
|  | Quadratic | 3 | 5 | 429.65 | -209.82 | 2 vs 3 | 0.07 | | 1 | | 0.799 |
|  | Null + Sex | 4 | 4 | 426.93 | -209.47 | 1 vs 4 | 1.02 | | 1 | | 0.313 |
| Neutral | **Null** | **1** | **3** | **470.08** | **-232.04** |  |  | |  | |  |
|  | Linear | 2 | 4 | 471.42 | -231.71 | 1 vs 2 | 0.65 | | 1 | | 0.418 |
|  | Quadratic | 3 | 5 | 472.72 | -231.36 | 2 vs 3 | 0.7 | | 1 | | 0.402 |
|  | Null + Sex | 4 | 4 | 472.07 | -232.03 | 1 vs 4 | 0.01 | | 1 | | 0.925 |
| **Testosterone (females)** | |  |  |  |  |  |  | |  | |  |
| *Left Amygdala* | |  |  |  |  |  |  | |  | |  |
| Angry | **Null** | **1** | **3** | **273.51** | **-133.75** |  |  | |  | |  |
|  | Linear | 2 | 4 | 275.49 | -133.75 | 1 vs 2 | 0.01 | | 1 | | 0.903 |
|  | Quadratic | 3 | 5 | 277.49 | -133.75 | 2 vs 3 | 0 | | 1 | | 0.997 |
| Fear | **Null** | **1** | **3** | **305.39** | **-149.69** |  |  | |  | |  |
|  | Linear | 2 | 4 | 306.14 | -149.07 | 1 vs 2 | 1.24 | | 1 | | 0.265 |
|  | Quadratic | 3 | 5 | 306.88 | -148.44 | 2 vs 3 | 1.26 | | 1 | | 0.262 |
| Sad | **Null** | **1** | **3** | **279.66** | **-136.83** |  |  | |  | |  |
|  | Linear | 2 | 4 | 281.34 | -136.67 | 1 vs 2 | 0.31 | | 1 | | 0.576 |
|  | Quadratic | 3 | 5 | 280.99 | -135.5 | 2 vs 3 | 2.35 | | 1 | | 0.125 |
| Happy | **Null** | **1** | **3** | **257.26** | **-125.63** |  |  | |  | |  |
|  | Linear | 2 | 4 | 259.07 | -125.54 | 1 vs 2 | 0.18 | | 1 | | 0.669 |
|  | Quadratic | 3 | 5 | 260.19 | -125.09 | 2 vs 3 | 0.89 | | 1 | | 0.346 |
| Neutral | **Null** | **1** | **3** | **263.38** | **-128.69** |  |  | |  | |  |
|  | Linear | 2 | 4 | 264.87 | -128.43 | 1 vs 2 | 0.51 | | 1 | | 0.474 |
|  | Quadratic | 3 | 5 | 264.38 | -127.19 | 2 vs 3 | 2.48 | | 1 | | 0.115 |
| *Right Amygdala* | |  |  |  |  |  |  | |  | |  |
| Angry | **Null** | **1** | **3** | **301.15** | **-147.58** |  |  | |  | |  |
|  | Linear | 2 | 4 | 302.1 | -147.05 | 1 vs 2 | 1.05 | | 1 | | 0.305 |
|  | Quadratic | 3 | 5 | 304.07 | -147.04 | 2 vs 3 | 0.03 | | 1 | | 0.863 |
| Fear | **Null** | **1** | **3** | **306.16** | **-150.08** |  |  | |  | |  |
|  | Linear | 2 | 4 | 308.02 | -150.01 | 1 vs 2 | 0.14 | | 1 | | 0.705 |
|  | Quadratic | 3 | 5 | 307.76 | -148.88 | 2 vs 3 | 2.26 | | 1 | | 0.133 |
| Sad | **Null** | **1** | **3** | **292.74** | **-143.37** |  |  | |  | |  |
|  | Linear | 2 | 4 | 294.72 | -143.36 | 1 vs 2 | 0.02 | | 1 | | 0.876 |
|  | Quadratic | 3 | 5 | 296.02 | -143.01 | 2 vs 3 | 0.7 | | 1 | | 0.402 |
| Happy | **Null** | **1** | **3** | **290.76** | **-142.38** |  |  | |  | |  |
|  | Linear | 2 | 4 | 291.71 | -141.86 | 1 vs 2 | 1.04 | | 1 | | 0.307 |
|  | Quadratic | 3 | 5 | 289.64 | -139.82 | 2 vs 3 | 4.07 | | 1 | | 0.044 |
| Neutral | **Null** | **1** | **3** | **298.62** | **-146.31** |  |  | |  | |  |
|  | Linear | 2 | 4 | 299.52 | -145.76 | 1 vs 2 | 1.1 | | 1 | | 0.294 |
|  | Quadratic | 3 | 5 | 300.87 | -145.43 | 2 vs 3 | 0.65 | | 1 | | 0.419 |
| *Left Nacc* |  |  |  |  |  |  |  | |  | |  |
| Angry | **Null** | **1** | **3** | **231.12** | **-112.56** |  |  | |  | |  |
|  | Linear | 2 | 4 | 231.68 | -111.84 | 1 vs 2 | 1.43 | | 1 | | 0.231 |
|  | Quadratic | 3 | 5 | 231.68 | -110.84 | 2 vs 3 | 2 | | 1 | | 0.157 |
| Fear | **Null** | **1** | **3** | **239.7** | **-116.85** |  |  | |  | |  |
|  | Linear | 2 | 4 | 241.43 | -116.71 | 1 vs 2 | 0.27 | | 1 | | 0.604 |
|  | Quadratic | 3 | 5 | 243.4 | -116.7 | 2 vs 3 | 0.03 | | 1 | | 0.87 |
| Sad | **Null** | **1** | **3** | **246.27** | **-120.13** |  |  | |  | |  |
|  | Linear | 2 | 4 | 248.19 | -120.09 | 1 vs 2 | 0.08 | | 1 | | 0.781 |
|  | Quadratic | 3 | 5 | 249.68 | -119.84 | 2 vs 3 | 0.51 | | 1 | | 0.475 |
| Happy | **Null** | **1** | **3** | **194.16** | **-94.08** |  |  | |  | |  |
|  | Linear | 2 | 4 | 195.86 | -93.93 | 1 vs 2 | 0.3 | | 1 | | 0.584 |
|  | Quadratic | 3 | 5 | 197.59 | -93.8 | 2 vs 3 | 0.27 | | 1 | | 0.602 |
| Neutral | **Null** | **1** | **3** | **241.86** | **-117.93** |  |  | |  | |  |
|  | Linear | 2 | 4 | 243.78 | -117.89 | 1 vs 2 | 0.08 | | 1 | | 0.781 |
|  | Quadratic | 3 | 5 | 245.78 | -117.89 | 2 vs 3 | 0 | | 1 | | 0.993 |
| *Right Nacc* |  |  |  |  |  |  |  | |  | |  |
| Angry | **Null** | **1** | **3** | **217.54** | **-105.77** |  |  | |  | |  |
|  | Linear | 2 | 4 | 219.42 | -105.71 | 1 vs 2 | 0.11 | | 1 | | 0.739 |
|  | Quadratic | 3 | 5 | 221.06 | -105.53 | 2 vs 3 | 0.37 | | 1 | | 0.544 |
| Fear | **Null** | **1** | **3** | **222.92** | **-108.46** |  |  | |  | |  |
|  | Linear | 2 | 4 | 224.36 | -108.18 | 1 vs 2 | 0.56 | | 1 | | 0.453 |
|  | Quadratic | 3 | 5 | 226.28 | -108.14 | 2 vs 3 | 0.08 | | 1 | | 0.78 |
| Sad | **Null** | **1** | **3** | **237.86** | **-115.93** |  |  | |  | |  |
|  | Linear | 2 | 4 | 238.81 | -115.4 | 1 vs 2 | 1.06 | | 1 | | 0.304 |
|  | Quadratic | 3 | 5 | 239.08 | -114.54 | 2 vs 3 | 1.73 | | 1 | | 0.189 |
| Happy | **Null** | **1** | **3** | **176.71** | **-85.35** |  |  | |  | |  |
|  | Linear | 2 | 4 | 177.41 | -84.7 | 1 vs 2 | 1.3 | | 1 | | 0.254 |
|  | Quadratic | 3 | 5 | 179.01 | -84.51 | 2 vs 3 | 0.39 | | 1 | | 0.53 |
| Neutral | **Null** | **1** | **3** | **221.98** | **-107.99** |  |  | |  | |  |
|  | Linear | 2 | 4 | 223.21 | -107.6 | 1 vs 2 | 0.77 | | 1 | | 0.379 |
|  | Quadratic | 3 | 5 | 224.94 | -107.47 | 2 vs 3 | 0.27 | | 1 | | 0.604 |
| *Left Hippocampus* | |  |  |  |  |  |  | |  | |  |
| Angry | **Null** | **1** | **3** | **214.13** | **-104.06** |  |  | |  | |  |
|  | Linear | 2 | 4 | 216.08 | -104.04 | 1 vs 2 | 0.05 | | 1 | | 0.826 |
|  | Quadratic | 3 | 5 | 218.01 | -104.01 | 2 vs 3 | 0.06 | | 1 | | 0.801 |
| Fear | **Null** | **1** | **3** | **253.65** | **-123.82** |  |  | |  | |  |
|  | Linear | 2 | 4 | 254.48 | -123.24 | 1 vs 2 | 1.17 | | 1 | | 0.28 |
|  | Quadratic | 3 | 5 | 254.77 | -122.38 | 2 vs 3 | 1.71 | | 1 | | 0.191 |
| Sad | **Null** | **1** | **3** | **214.96** | **-104.48** |  |  | |  | |  |
|  | Linear | 2 | 4 | 216.94 | -104.47 | 1 vs 2 | 0.02 | | 1 | | 0.901 |
|  | Quadratic | 3 | 5 | 217.7 | -103.85 | 2 vs 3 | 1.24 | | 1 | | 0.265 |
| Happy | **Null** | **1** | **3** | **196.3** | **-95.15** |  |  | |  | |  |
|  | Linear | 2 | 4 | 196.7 | -94.35 | 1 vs 2 | 1.61 | | 1 | | 0.205 |
|  | Quadratic | 3 | 5 | 198.63 | -94.32 | 2 vs 3 | 0.06 | | 1 | | 0.802 |
| Neutral | **Null** | **1** | **3** | **212.02** | **-103.01** |  |  | |  | |  |
|  | Linear | 2 | 4 | 213.4 | -102.7 | 1 vs 2 | 0.61 | | 1 | | 0.434 |
|  | Quadratic | 3 | 5 | 214.52 | -102.26 | 2 vs 3 | 0.89 | | 1 | | 0.346 |
| *Right Hippocampus* | |  |  |  |  |  |  | |  | |  |
| Angry | **Null** | **1** | **3** | **246.01** | **-120** |  |  | |  | |  |
|  | Linear | 2 | 4 | 247.07 | -119.53 | 1 vs 2 | 0.94 | | 1 | | 0.333 |
|  | Quadratic | 3 | 5 | 249.03 | -119.51 | 2 vs 3 | 0.04 | | 1 | | 0.836 |
| Fear | **Null** | **1** | **3** | **238.66** | **-116.33** |  |  | |  | |  |
|  | Linear | 2 | 4 | 240.56 | -116.28 | 1 vs 2 | 0.1 | | 1 | | 0.757 |
|  | Quadratic | 3 | 5 | 238.85 | -114.43 | 2 vs 3 | 3.7 | | 1 | | 0.054 |
| Sad | **Null** | **1** | **3** | **217.5** | **-105.75** |  |  | |  | |  |
|  | Linear | 2 | 4 | 219.43 | -105.71 | 1 vs 2 | 0.07 | | 1 | | 0.791 |
|  | Quadratic | 3 | 5 | 219.33 | -104.67 | 2 vs 3 | 2.1 | | 1 | | 0.148 |
| Happy | **Null** | **1** | **3** | **208.58** | **-101.29** |  |  | |  | |  |
|  | Linear | 2 | 4 | 207.89 | -99.95 | 1 vs 2 | 2.68 | | 1 | | 0.101 |
|  | Quadratic | 3 | 5 | 207.11 | -98.56 | 2 vs 3 | 2.78 | | 1 | | 0.095 |
| Neutral | **Null** | **1** | **3** | **233.52** | **-113.76** |  |  | |  | |  |
|  | Linear | 2 | 4 | 233.58 | -112.79 | 1 vs 2 | 1.94 | | 1 | | 0.164 |
|  | Quadratic | 3 | 5 | 235.49 | -112.75 | 2 vs 3 | 0.09 | | 1 | | 0.766 |
| **Testosterone (males)** | |  |  |  |  |  |  | |  | |  |
| *Left Amygdala* | |  |  |  |  |  |  | |  | |  |
| Angry | **Null** | **1** | **3** | **282.42** | **-138.21** |  |  | |  | |  |
|  | Linear | 2 | 4 | 284.07 | -138.04 | 1 vs 2 | 0.35 | | 1 | | 0.553 |
|  | Quadratic | 3 | 5 | 285.38 | -137.69 | 2 vs 3 | 0.69 | | 1 | | 0.407 |
| Fear | **Null** | **1** | **3** | **269.34** | **-131.67** |  |  | |  | |  |
|  | Linear | 2 | 4 | 271.22 | -131.61 | 1 vs 2 | 0.12 | | 1 | | 0.724 |
|  | Quadratic | 3 | 5 | 273 | -131.5 | 2 vs 3 | 0.22 | | 1 | | 0.636 |
| Sad | **Null** | **1** | **3** | **280.08** | **-137.04** |  |  | |  | |  |
|  | Linear | 2 | 4 | 282.07 | -137.04 | 1 vs 2 | 0.01 | | 1 | | 0.91 |
|  | Quadratic | 3 | 5 | 284.01 | -137.01 | 2 vs 3 | 0.06 | | 1 | | 0.807 |
| Happy | **Null** | **1** | **3** | **276.59** | **-135.29** |  |  | |  | |  |
|  | Linear | 2 | 4 | 278.39 | -135.2 | 1 vs 2 | 0.2 | | 1 | | 0.656 |
|  | Quadratic | 3 | 5 | 280.27 | -135.14 | 2 vs 3 | 0.12 | | 1 | | 0.733 |
| Neutral | **Null** | **1** | **3** | **254.74** | **-124.37** |  |  | |  | |  |
|  | Linear | 2 | 4 | 256.72 | -124.36 | 1 vs 2 | 0.02 | | 1 | | 0.876 |
|  | Quadratic | 3 | 5 | 258.65 | -124.33 | 2 vs 3 | 0.06 | | 1 | | 0.8 |
| *Right Amygdala* | |  |  |  |  |  |  | |  | |  |
| Angry | **Null** | **1** | **3** | **277.07** | **-135.54** |  |  | |  | |  |
|  | Linear | 2 | 4 | 278.33 | -135.17 | 1 vs 2 | 0.74 | | 1 | | 0.39 |
|  | Quadratic | 3 | 5 | 279.02 | -134.51 | 2 vs 3 | 1.31 | | 1 | | 0.253 |
| Fear | **Null** | **1** | **3** | **245.15** | **-119.57** |  |  | |  | |  |
|  | Linear | 2 | 4 | 246.88 | -119.44 | 1 vs 2 | 0.27 | | 1 | | 0.603 |
|  | Quadratic | 3 | 5 | 248.44 | -119.22 | 2 vs 3 | 0.44 | | 1 | | 0.509 |
| Sad | **Null** | **1** | **3** | **254.03** | **-124.01** |  |  | |  | |  |
|  | Linear | 2 | 4 | 256.01 | -124.01 | 1 vs 2 | 0.02 | | 1 | | 0.897 |
|  | Quadratic | 3 | 5 | 255.64 | -122.82 | 2 vs 3 | 2.37 | | 1 | | 0.124 |
| Happy | **Null** | **1** | **3** | **262.01** | **-128.01** |  |  | |  | |  |
|  | Linear | 2 | 4 | 261.35 | -126.67 | 1 vs 2 | 2.66 | | 1 | | 0.103 |
|  | Quadratic | 3 | 5 | 262.91 | -126.46 | 2 vs 3 | 0.43 | | 1 | | 0.511 |
| Neutral | **Null** | **1** | **3** | **261.94** | **-127.97** |  |  | |  | |  |
|  | Linear | 2 | 4 | 263.3 | -127.65 | 1 vs 2 | 0.64 | | 1 | | 0.423 |
|  | Quadratic | 3 | 5 | 265.21 | -127.61 | 2 vs 3 | 0.09 | | 1 | | 0.762 |
| *Left Nacc* |  |  |  |  |  |  |  | |  | |  |
| Angry | **Null** | **1** | **3** | **247.46** | **-120.73** |  |  | |  | |  |
|  | Linear | 2 | 4 | 249.23 | -120.61 | 1 vs 2 | 0.24 | | 1 | | 0.627 |
|  | Quadratic | 3 | 5 | 250.4 | -120.2 | 2 vs 3 | 0.82 | | 1 | | 0.364 |
| Fear | **Null** | **1** | **3** | **239.82** | **-116.91** |  |  | |  | |  |
|  | Linear | 2 | 4 | 241.54 | -116.77 | 1 vs 2 | 0.28 | | 1 | | 0.597 |
|  | Quadratic | 3 | 5 | 243.32 | -116.66 | 2 vs 3 | 0.21 | | 1 | | 0.644 |
| Sad | **Null** | **1** | **3** | **249.18** | **-121.59** |  |  | |  | |  |
|  | Linear | 2 | 4 | 251.11 | -121.56 | 1 vs 2 | 0.07 | | 1 | | 0.789 |
|  | Quadratic | 3 | 5 | 252.85 | -121.42 | 2 vs 3 | 0.27 | | 1 | | 0.606 |
| Happy | **Null** | **1** | **3** | **260.51** | **-127.26** |  |  | |  | |  |
|  | Linear | 2 | 4 | 262.09 | -127.04 | 1 vs 2 | 0.42 | | 1 | | 0.515 |
|  | Quadratic | 3 | 5 | 264.04 | -127.02 | 2 vs 3 | 0.05 | | 1 | | 0.828 |
| Neutral | **Null** | **1** | **3** | **261.91** | **-127.96** |  |  | |  | |  |
|  | Linear | 2 | 4 | 263.38 | -127.69 | 1 vs 2 | 0.53 | | 1 | | 0.465 |
|  | Quadratic | 3 | 5 | 265.36 | -127.68 | 2 vs 3 | 0.01 | | 1 | | 0.909 |
| *Right Nacc* |  |  |  |  |  |  |  | |  | |  |
| Angry | **Null** | **1** | **3** | **244.24** | **-119.12** |  |  | |  | |  |
|  | Linear | 2 | 4 | 245.47 | -118.74 | 1 vs 2 | 0.77 | | 1 | | 0.382 |
|  | Quadratic | 3 | 5 | 247.17 | -118.59 | 2 vs 3 | 0.3 | | 1 | | 0.586 |
| Fear | **Null** | **1** | **3** | **225.23** | **-109.61** |  |  | |  | |  |
|  | Linear | 2 | 4 | 226.8 | -109.4 | 1 vs 2 | 0.42 | | 1 | | 0.517 |
|  | Quadratic | 3 | 5 | 228.6 | -109.3 | 2 vs 3 | 0.21 | | 1 | | 0.647 |
| Sad | **Null** | **1** | **3** | **259.34** | **-126.67** |  |  | |  | |  |
|  | Linear | 2 | 4 | 260.92 | -126.46 | 1 vs 2 | 0.41 | | 1 | | 0.52 |
|  | Quadratic | 3 | 5 | 262.84 | -126.42 | 2 vs 3 | 0.08 | | 1 | | 0.781 |
| Happy | **Null** | **1** | **3** | **250.75** | **-122.37** |  |  | |  | |  |
|  | Linear | 2 | 4 | 252.74 | -122.37 | 1 vs 2 | 0.01 | | 1 | | 0.936 |
|  | Quadratic | 3 | 5 | 254.71 | -122.35 | 2 vs 3 | 0.03 | | 1 | | 0.864 |
| Neutral | **Null** | **1** | **3** | **253.83** | **-123.92** |  |  | |  | |  |
|  | Linear | 2 | 4 | 255.06 | -123.53 | 1 vs 2 | 0.77 | | 1 | | 0.38 |
|  | Quadratic | 3 | 5 | 256.72 | -123.36 | 2 vs 3 | 0.35 | | 1 | | 0.556 |
| *Left Hippocampus* | |  |  |  |  |  |  | |  | |  |
| Angry | **Null** | **1** | **3** | **228.32** | **-111.16** |  |  | |  | |  |
|  | Linear | 2 | 4 | 228.8 | -110.4 | 1 vs 2 | 1.52 | | 1 | | 0.217 |
|  | Quadratic | 3 | 5 | 227.35 | -108.68 | 2 vs 3 | 3.44 | | 1 | | 0.063 |
| Fear | **Null** | **1** | **3** | **230.9** | **-112.45** |  |  | |  | |  |
|  | Linear | 2 | 4 | 232.04 | -112.02 | 1 vs 2 | 0.86 | | 1 | | 0.352 |
|  | Quadratic | 3 | 5 | 233.97 | -111.99 | 2 vs 3 | 0.06 | | 1 | | 0.806 |
| Sad | **Null** | **1** | **3** | **232.66** | **-113.33** |  |  | |  | |  |
|  | Linear | 2 | 4 | 234.29 | -113.14 | 1 vs 2 | 0.37 | | 1 | | 0.542 |
|  | Quadratic | 3 | 5 | 235.95 | -112.97 | 2 vs 3 | 0.34 | | 1 | | 0.559 |
| Happy | **Null** | **1** | **3** | **243.26** | **-118.63** |  |  | |  | |  |
|  | Linear | 2 | 4 | 245.24 | -118.62 | 1 vs 2 | 0.02 | | 1 | | 0.882 |
|  | Quadratic | 3 | 5 | 247.24 | -118.62 | 2 vs 3 | 0 | | 1 | | 0.946 |
| Neutral | **Null** | **1** | **3** | **230.69** | **-112.35** |  |  | |  | |  |
|  | Linear | 2 | 4 | 232.69 | -112.35 | 1 vs 2 | 0 | | 1 | | 0.951 |
|  | Quadratic | 3 | 5 | 234.69 | -112.34 | 2 vs 3 | 0 | | 1 | | 0.961 |
| *Right Hippocampus* | |  |  |  |  |  |  | |  | |  |
| Angry | **Null** | **1** | **3** | **243.92** | **-118.96** |  |  | |  | |  |
|  | Linear | 2 | 4 | 244.3 | -118.15 | 1 vs 2 | 1.62 | | 1 | | 0.203 |
|  | Quadratic | 3 | 5 | 245.17 | -117.59 | 2 vs 3 | 1.12 | | 1 | | 0.289 |
| Fear | **Null** | **1** | **3** | **221** | **-107.5** |  |  | |  | |  |
|  | Linear | 2 | 4 | 222.5 | -107.25 | 1 vs 2 | 0.5 | | 1 | | 0.479 |
|  | Quadratic | 3 | 5 | 224.47 | -107.23 | 2 vs 3 | 0.03 | | 1 | | 0.852 |
| Sad | **Null** | **1** | **3** | **219.19** | **-106.59** |  |  | |  | |  |
|  | Linear | 2 | 4 | 221.19 | -106.59 | 1 vs 2 | 0 | | 1 | | 0.997 |
|  | Quadratic | 3 | 5 | 222.55 | -106.27 | 2 vs 3 | 0.64 | | 1 | | 0.423 |
| Happy | **Null** | **1** | **3** | **220.24** | **-107.12** |  |  | |  | |  |
|  | Linear | 2 | 4 | 222.11 | -107.05 | 1 vs 2 | 0.14 | | 1 | | 0.713 |
|  | Quadratic | 3 | 5 | 223.91 | -106.95 | 2 vs 3 | 0.2 | | 1 | | 0.653 |
| Neutral | **Null** | **1** | **3** | **241.02** | **-117.51** |  |  | |  | |  |
|  | Linear | 2 | 4 | 242.6 | -117.3 | 1 vs 2 | 0.42 | | 1 | | 0.517 |
|  | Quadratic | 3 | 5 | 244.57 | -117.29 | 2 vs 3 | 0.03 | | 1 | | 0.87 |

**Table S8. Whole brain results for “age” model.**

| **Cluster** | **Region** | **Hemi** | | **Cluster Size** | **Z Stat** | **X** | | **Y** | | **Z** |
| --- | --- | --- | --- | --- | --- | --- | --- | --- | --- | --- |
| **Age** |  | |  |  |  |  | |  | |  |
| 1 | Cuneus/Superior Occipital Cortex | | R | 115 | 7.7107 | 27 | | -88 | | 32 |
| 2 | Inferior Occipital Cortex | | R | 72 | 7.5961 | 45 | | -82 | | 2 |
| 3 | Inferior Occipital Cortex | | L | 38 | 7.0499 | -39 | | -88 | | -7 |
| 4 | Superior Occipital Cortex | | L | 35 | 4.5499 | -27 | | -88 | | 32 |
| **Age*age** |  | |  |  |  |  | |  | |  |
| 1 | Cuneus/Lateral Occipital Cortex | | R | 943 | -7.59 | 42 | | -91 | | 14 |
| 2 | Cuneus/Lateral Occipital Cortex | | L | 341 | -7.1043 | -12 | | -100 | | 26 |
| 3 | Cerebellum | | R | 195 | -5.1608 | 9 | | -34 | | -52 |
| 4 | Cerebellum | | L | 172 | -6.4809 | -36 | | -70 | | -25 |
| 5 | Mid Cingulate (WM) | | R | 129 | -6.1766 | 24 | | -22 | | 23 |
| 6 | Somatosensory Cortex | | R | 126 | 5.2421 | 21 | | -31 | | 83 |
| 7 | anterior-rostral PFC | |  | 122 | -6.1261 | 12 | | 74 | | -7 |
| 8 | Primary Motor Cortex | | R | 101 | 4.2996 | 36 | | -22 | | 59 |
| 9 | vmPFC/mOFC | | L | 92 | -5.1727 | -9 | | 53 | | -22 |
| 10 | Parahippocampus | | L | 77 | -5.2398 | -24 | | -34 | | -4 |
| 11 | vlPFC | | L | 76 | -4.4066 | -27 | | 50 | | 2 |
| 12 | Mid Cingulate (WM) | | L | 75 | -6.4636 | -24 | | -19 | | 29 |
| 13 | Superior parietal | | L | 70 | -5.7156 | -27 | | -61 | | 59 |
| 14 | Primary Motor Cortex | | L | 70 | 4.9595 | -12 | | -13 | | 77 |
| 15 | Brainstem | | R | 57 | -6.4661 | 12 | | -4 | | -28 |
| 16 | Premotor Cortex | | R | 51 | 4.8738 | 24 | | 5 | | 62 |
| 17 | Lingual Gyrus | | L | 47 | 4.8576 | -15 | | -55 | | -1 |
| 18 | dlPFC | | L | 39 | -4.3156 | -21 | | 29 | | 62 |
| 19 | pSTS | | R | 37 | 4.1372 | 60 | | -43 | | 11 |
| 20 | Primary Motor Cortex | | R | 37 | 5.9245 | 42 | | -31 | | 68 |
| 21 | mOFC | | R | 35 | -4.6617 | 12 | | 23 | | -1 |
| 22 | Somatosensory Cortex | | L | 34 | -4.636 | -57 | | -13 | | 26 |
| **Age*age*sex** |  | |  |  |  |  |  | |  |  |
| 1 | Cuneus / Later Occipital Cortex | |  | 2033 | -9.3584 | 3 | | -94 | | 8 |
| 2 | Motor Cortices | | R | 676 | -6.9145 | 39 | | -31 | | 50 |
| 3 | vACC/vmPFC | |  | 630 | 6.1479 | 6 | | 41 | | 2 |
| 4 | Primary MotorCortex | | L | 354 | -6.4801 | -27 | | -22 | | 80 |
| 5 | Inferior Occipital Cortex | | L | 136 | -6.79 | -45 | | -79 | | -19 |
| 6 | Mid Cingulate/dACC | | L | 92 | 6.5923 | -12 | | -1 | | 26 |
| 7 | OFC | | R | 69 | -4.5571 | 18 | | 32 | | -25 |
| 8 | Anterior Temporal Cortex | | R | 58 | 5.282 | 48 | | -4 | | -37 |
| 9 | Cerebellum/Fusiform | | L | 48 | -4.6419 | -33 | | -46 | | -25 |
| 10 | Cerebellum | | L | 46 | 5.3354 | -30 | | -73 | | -49 |
| 11 | Parahippocampus | | R | 45 | 5.0432 | 18 | | -16 | | -22 |
| 12 | Superior Parietal Cortex | | L | 44 | -5.928 | -36 | | -49 | | 71 |
| 13 | Inferior Temporal Cortex | | L | 35 | 4.9338 | -48 | | -37 | | -16 |
| 14 | Insula | | L | 35 | 5.7195 | -39 | | 5 | | 14 |
| 15 | Frontal Eye Field (WM) | | L | 34 | 4.524 | -33 | | 2 | | 35 |

**Table S9. Whole brain results for “PDS” model**

| **Cluster** | **Region** | **Hemi** | **Cluster Size** | **Z Stat** | **X** | **Y** | **Z** |
| --- | --- | --- | --- | --- | --- | --- | --- |
| **PDS** |  |  |  |  |  |  |  |
| 1 | Cuneus/Lateral Occipital Cortex |  | 629 | 6.9556 | 27 | -85 | 29 |
| 2 | Inferior Occipital Cortex | L | 117 | 5.9135 | -36 | -91 | -7 |
| 3 | R mOFC | R | 60 | 4.441 | 9 | 20 | -22 |
| 4 | Primary Motor Cortex | L | 53 | 5.2553 | -42 | -28 | 68 |
| **PDS*PDS** | |  |  |  |  |  |  |
| 1 | Cuneus |  | 462 | 6.6686 | -15 | -79 | 5 |
| 2 | Somatosensory Motor Cortex | R | 89 | 5.4628 | 48 | -37 | 68 |
| 3 | SMA |  | 67 | 4.7201 | 0 | -22 | 80 |
| 4 | Inferior Occipital Cortex/Fusiform | L | 56 | 7.2708 | -45 | -85 | -13 |
| 5 | Lingual Gyrus | R | 52 | 4.692 | 18 | -73 | -7 |
| 6 | Premotor Cortex | R | 52 | 5.3463 | 30 | -1 | 74 |
| 7 | Angular Gyrus/ TPJ | L | 48 | -4.6999 | -66 | -46 | 38 |
| 8 | Hippocampus / Amygdala | R | 38 | -4.3809 | 24 | -13 | -16 |
| 9 | Lingual Gyrus | R | 34 | 4.1632 | 24 | -94 | -10 |
| 10 | Mid Cingulate |  | 33 | 4.9056 | 6 | -13 | 29 |
| **PDS*PDS*sex** | |  |  |  |  |  |  |
| 1 | vAcc / vMPFC |  | 269 | 6.0965 | 12 | 38 | -1 |
| 2 | Cuneus / Lateral Occipital Cortex |  | 194 | -6.2903 | 3 | -94 | 8 |
| 3 | dmPFC | R | 116 | 5.4888 | 12 | 59 | 14 |
| 4 | Inferior Occipital Cortex / Fusiform | L | 103 | -7.3426 | -45 | -82 | -16 |
| 5 | Lingual Gyrus / Fusiform | R | 92 | -5.0639 | 30 | -64 | -16 |
| 6 | Caudate / WM | R | 58 | 4.9066 | 18 | 23 | -4 |
| 7 | Cerebellum | L | 47 | -5.8061 | -30 | -52 | -25 |
| 8 | Cerebellum | L | 45 | -5.8993 | -54 | -64 | -34 |
| 9 | Intraparietal Sulcus | R | 44 | -4.3811 | 36 | -55 | 35 |
| 10 | Precuneus | R | 41 | 4.4506 | 6 | -61 | 29 |
| 11 | Precuneus / Cuneus | L | 40 | 4.302 | -12 | -73 | 23 |
| 12 | Premotor Cortex | R | 39 | -4.4382 | 54 | 5 | 38 |
| 13 | NAcc | R | 36 | 4.8931 | 12 | 5 | -4 |
| 14 | OFC | L | 33 | 4.574 | -33 | 32 | -10 |
